# Heat Shock Factor 1 Governs Sleep-Wake Cycles Across Species

**DOI:** 10.1101/2024.11.15.623879

**Authors:** Shintaro Yamazaki, Utham K. Valekunja, Jing Chen-Roetling, Akhilesh B. Reddy

## Abstract

Heat Shock Factor 1 (HSF1) is a critical transcription factor for cellular proteostasis, but its role in sleep regulation remains unexplored. We demonstrate that nuclear HSF1 levels in the mouse brain fluctuate with sleep-wake cycles, increasing during extended wakefulness and decreasing during sleep. Using CUT&RUN and RNA-seq, we identified HSF1-regulated transcriptional changes involved in synaptic organization, expanding its known functions beyond traditional heat shock responses. Both systemic and brain-specific *Hsf1* knockout mice exhibit altered sleep homeostasis, including increased delta power after sleep deprivation and upregulation of sleep-related genes. However, these knockouts struggle to maintain sleep due to disrupted synaptic organization. In *Drosophila*, knockout of HSF1’s ortholog results in fragmented sleep patterns, suggesting a conserved role for HSF1 in sleep regulation across species. Our findings reveal a novel molecular mechanism underlying sleep regulation and offer potential therapeutic targets for sleep disturbances.

## Introduction

Sleep is a fundamental biological process conserved across species, playing a crucial role in maintaining cognitive function, emotional stability, and overall health ^1–6^. Despite its importance, the molecular mechanisms governing sleep regulation remain incompletely understood. Uncovering key regulatory molecules involved in sleep maintenance is essential for comprehending both normal sleep physiology and the pathophysiology of sleep disorders.

Heat Shock Factor 1 (HSF1), a master transcription factor primarily known for its role in cellular stress responses, particularly in regulating proteostasis through the induction of heat shock proteins (HSPs) ^7,8^, has been extensively studied in the context of cancer and neurodegenerative diseases ^9–11^. However, its potential involvement in sleep regulation remains unexplored. Recent observations have highlighted the dynamic nature of gene expression and protein regulation during sleep-wake cycles, suggesting that transcriptional regulators like HSF1 may play vital roles in sleep homeostasis ^12–14^.

The cerebral cortex plays a prominent role in sleep-wake processes. Cortical neurons exhibit distinct firing patterns during different sleep stages, and their activity is tightly linked to sleep homeostasis ^15^. The cortex undergoes significant changes in gene expression and synaptic plasticity during sleep and wake states ^16,17^. Moreover, slow-wave activity in the cortex, a key marker of sleep pressure, increases with prolonged wakefulness and decreases during sleep, making it an ideal site for investigating molecular mechanisms of sleep homeostasis ^18^. Recent studies have also highlighted the importance of cortical glial cells in sleep regulation, further emphasizing the cortex’s significance in sleep ^19^.

In this study, we investigate the role of HSF1 in sleep regulation using a multi-faceted approach. We examine HSF1 expression patterns in the brain across sleep-wake cycles and under conditions of sleep deprivation. Using conditional knockout mouse models and *Drosophila* mutants, we assess the impact of HSF1 deficiency on sleep architecture, including non-rapid eye movement (NREM) sleep, rapid eye movement (REM) sleep, and wakefulness patterns. Additionally, we employ CUT&RUN and RNA sequencing to identify HSF1-regulated genes and pathways, delineating the molecular mechanisms underlying HSF1’s influence on sleep.

Our findings reveal a novel role for HSF1 in maintaining sleep continuity and quality, independent of its well-established functions in stress responses. We also uncover a previously unrecognized function of HSF1 in regulating synaptic organization, which appears to be critical for sustaining sleep. This study not only expands our understanding of HSF1’s biological functions but also provides new insights into the molecular underpinnings of sleep regulation. By elucidating HSF1’s role in sleep, we open new avenues for investigating sleep disorders and potentially developing targeted therapeutic interventions to improve sleep quality and overall health.

## Results

### HSF1 nuclear expression in the brain oscillates with sleep-wake cycles

Previous studies have shown that HSF1 oscillate between the nucleus and cytoplasm in peripheral organs including liver tissue ^20^. Consistent with these findings, our data indicate that nuclear HSF1 expression in the brain exhibited a diurnal pattern, increasing during wakefulness and decreasing during sleep. Immunoblot analysis of nuclear and total cellular protein in the cerebral cortex of C57BL/6J mice (n = 3 per time point) at 4-hour intervals over a 24-hour period revealed significant fluctuations in nuclear HSF1 levels, while total cellular HSF1 remained constant during the 12h:12h light-dark (LD) cycle (Fig. 1a-c). Quantitative PCR (qPCR) of the same brain tissues showed constant *Hsf1* mRNA expression throughout the cycle (Fig. 1d), whereas circadian genes *Per2* and *Bmal1* displayed rhythmic expression patterns (Fig. 1e,f). These results suggest that HSF1 translocates between the cytoplasm and nucleus, with nuclear accumulation occurring during the active phase (ZT12-24) and cytoplasmic localization during the inactive phase (ZT0-12).

**Figure 1:**
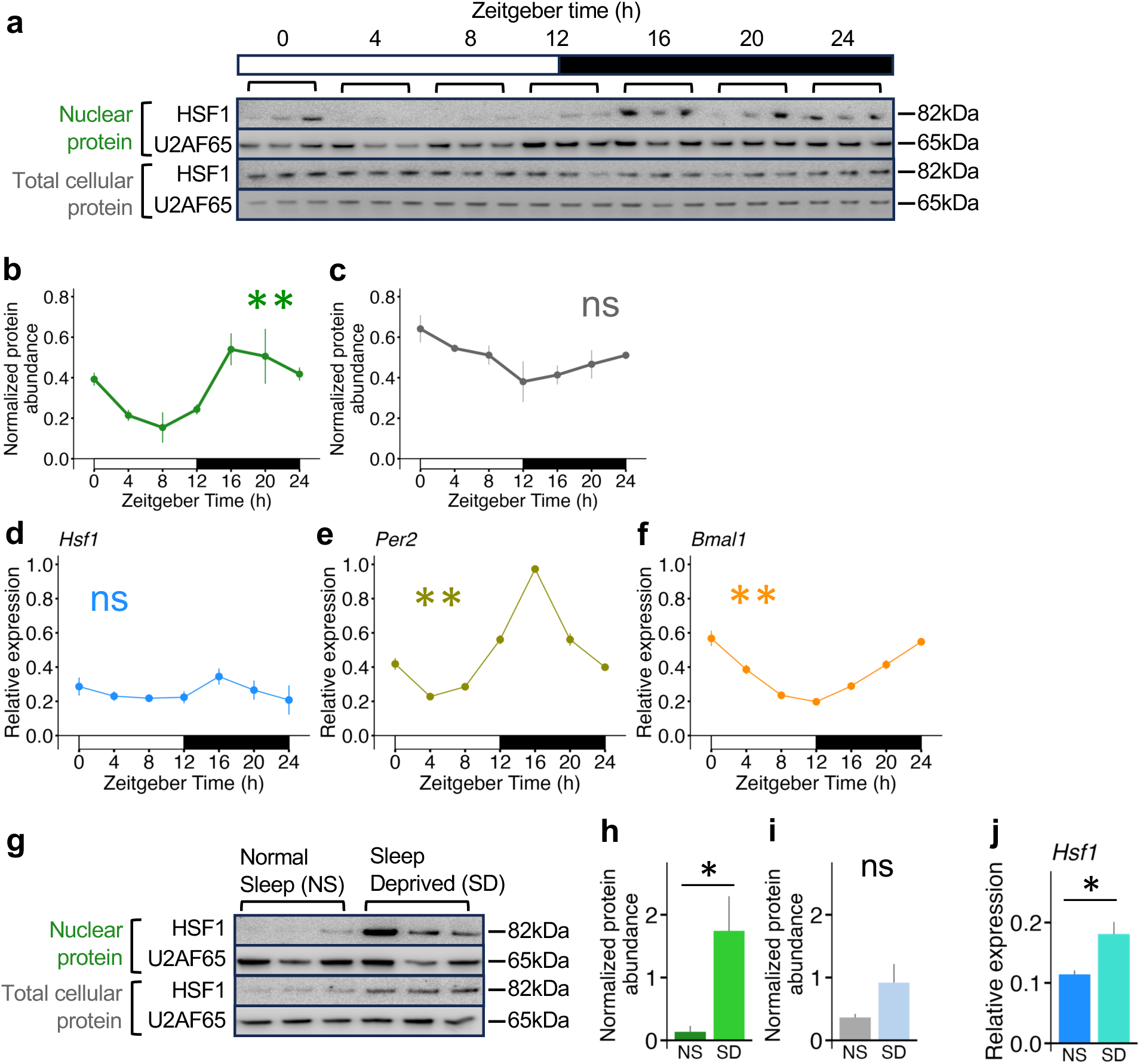
Nuclear HSF1 expression in mouse brain increase with extended wakefulness, while decrease during sleep. **(a)** Immunoblot showing the expressions of HSF1 and U2AF65 (loading control for nuclear protein) in nuclear and total cellular protein extracted from cerebral cortex tissues of C57BL/6J mice (n=3 per time point) at each time point: every four hours (ZT0, 4, 8, 12, 16, 20 and 24). (**b), (c)** Line plots showing the immunoblot band intensities of HSF1 normalized to U2AF65 at each time point for nuclear protein (b) and total cellular protein (c). **(d), (e), (f)** Line plots showing the mRNA expression of *Hsf1* (d)*, Per2* (e) *or Bmal1* (f) normalized to *Actb* at each time point (n=3 per time point). **(g)** Immunoblot showing the expressions of HSF1 and U2AF65 in nuclear and total cellular protein extracted from cerebral cortex tissues of C57BL/6J mice (n=3 per group) for normal sleep (NS) and sleep deprived (SD) group (n=3 per group) at ZT6. **(h), (i)** Bar plots showing the immunoblot band intensities of HSF1 normalized to U2AF65 in nuclear (h) or total cellular protein (i) comparing NS and SD group. **(j)** Bar plot showing the mRNA expression of *Hsf1* normalized to *Actb* comparing NS and SD group (n=3 per group). Data are mean ±s.e.m. Statistical significance was determined using one-way ANOVA (b,c,d,e,f), two-tailed t-test (h,i,j). *: p<0.05, **: p<0.01, ns: not significant.

To further investigate the relationship between nuclear HSF1 expression and sleep/wake states, we compared brain tissues from mice subjected to 6 hours of sleep deprivation (SD, ZT0-6) with those from normal sleep (NS) mice. Immunoblotting showed that extended wakefulness induced by SD significantly upregulated nuclear HSF1 expression (Fig. 1g-i) and *Hsf1* mRNA levels (Fig. 1j), indicating a causal relationship between HSF1 expression and sleep/wake states. Collectively, these findings demonstrate the dynamic regulation of HSF1 expression in the brain in response to sleep and wakefulness, implicating HSF1 as a critical player in the molecular mechanisms underlying sleep regulation.

#### Hsf1 knockouts have reduced NREM sleep despite persistent sleep pressure

To investigate HSF1’s functional role in sleep regulation, we established two conditional *Hsf1* knockout mouse models using the Cre-LoxP system: *CAG^CreERTM^;Hsf1^flox/flox^* for systemic *Hsf1* knockout upon tamoxifen injection, and *Hsf1^flox/flox^* for central nervous system (CNS)-specific knockout via retro-orbital sinus delivery of adeno-associated virus (AAV)-CAG-Cre-GFP (PHP.eB) (Fig. 2a,b, Supplementary Fig. 1). Sleep analysis in both systemic and brain-specific *Hsf1* knockout models revealed significant reductions in non-REM (NREM) sleep accompanied by increased wakefulness, particularly during the light phase (ZT0-12) and over the entire 24-hour period (Fig. 2c-e, i-k).

**Figure 2:**
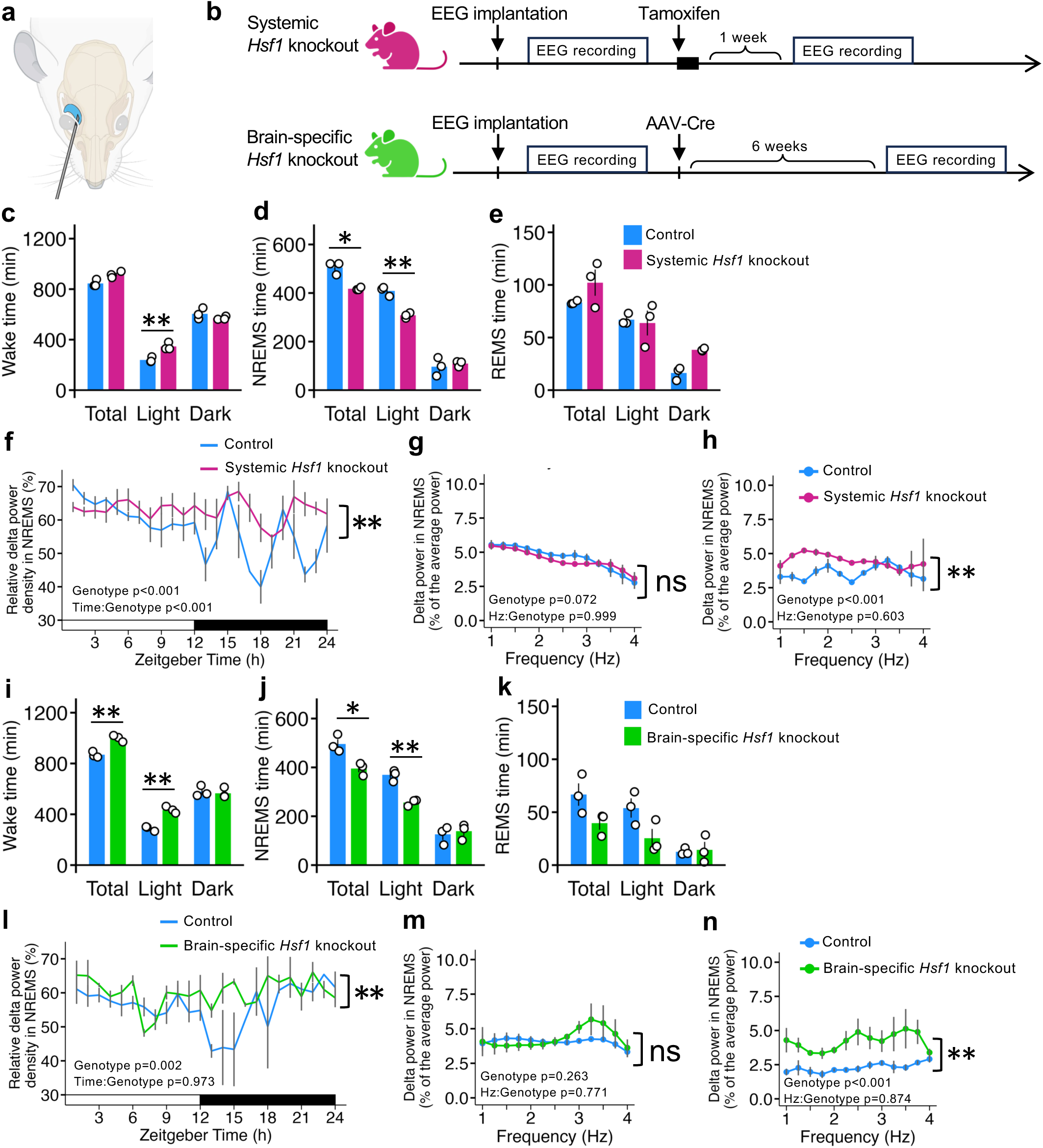
Hsf1 knockout mice reduce NREM sleep and increase wakefulness despite the sleep demands. **(a)** Schematic showing the AAV delivery rout via retro-orbital sinus. **(b)** Schematic showing the experiment time courses for the conditional *Hsf1* knockout models. Systemic knockout model: *CAG^CreERTM^;Hsf1^flox/flox^*administered corn oil (Control), or tamoxifen (Systemic *Hsf1* knockout), brain-specific knockout model: *Hsf1^flox/flox^* administered AAV-CAG-GFP (Control), or AAV-CAG-Cre-GFP (Brain-specific *Hsf1* knockout). n=3 per group. **(c), (d), (e), (i), (j), (k)** Bar plots showing the total time (minutes) spent in Wake (c,i), NREM sleep (d,j) or REM sleep (e,k) in each phase (Total: ZT0-24, Light: ZT0-12 or Dark: ZT12-24) for the systemic (c,d,e) and the brain-specific *Hsf1* knockout model (i,j,k). **(f), (l)** Line plots showing the hourly average of the delta power (1-4Hz) density in NREM sleep (%) for the systemic (f) and the brain-specific *Hsf1* knockout model (l). **(g), (h), (m), (n)** Line plots showing the average delta power density in NREM sleep (%) during the entire 24h period (ZT0-24, g,m) or 2 hours after light phase (ZT12-14, h,n) for the systemic (g,h) and the brain-specific *Hsf1* knockout model (m,n), comparing Control and *Hsf1* knockout group in 1-4Hz range. Data are mean ±s.e.m. Statistical significance was determined using two-way ANOVA with Tukey’s HSD (c,d,e,f,g,h,i,j,k,l,m,n). *: p<0.05, **: p<0.01, ns: not significant. P values for main effect of genotype factor: control or *Hsf1* knockout (Genotype), and interaction effect between genotype and time factors: every hour from ZT1 to ZT24 (Time:Genotype) (f,g,h,l,m,n).

Although the increased wake time was accompanied partially by elevated locomotor activity during baseline period (Supplementary Fig. 2d,e,p,q), correlation analysis revealed that sleep stage transitions towards wake, such as NREM sleep-to-wake (NtoW) or REM sleep-to-wake (RtoW), did not directly lead to significant increases in locomotor activity in either systemic or brain-specific *Hsf1* knockout mice (Supplementary Fig. 2y). While the knockouts had increased wake time, they exhibited quiescent behavior after waking, which suggests that the disrupted sleep-wake pattern in *Hsf1* knockout mice is not driven by increased activity but rather reflects alterations in sleep regulatory process ^21^.

The pattern of hourly relative delta power density during NREM sleep, an indicator of sleep pressure ^22,23^, differed markedly between genotypes. *Hsf1* knockout mice failed to decrease delta power density after the light phase, during which they slept less compared to controls (Fig. 2f,l). While no significant difference in delta power was seen over the entire 24-hour period (ZT0-24) between knockout and control mice (Fig. 2g,m), knockout mice did not exhibit the expected decrease in delta power during ZT12-14, the period immediately following the inactive phase (Fig. 2h,n). These findings suggest that the loss of *Hsf1* reduces sleep quantity while maintaining normal sleep pressure, as evidenced by the persistent delta power. Despite consistent sleep demand, *Hsf1* knockout mice did not increase sleep during the light-dark cycle, implying a role for HSF1 in initiating or maintaining the sleep state.

#### Hsf1 knockout mice exhibit fragmented sleep

Analysis of sleep-wake architecture revealed markedly fragmented sleep patterns in *Hsf1* knockout mice. Systemic *Hsf1* knockout mice showed an increased numbers of wake and sleep bouts, a trend also observed in brain-specific knockouts (Fig. 3a,b,g,h). However, the mean duration of these sleep bouts was reduced in both knockout models (Fig. 3c,i). Further analysis of sleep bout characteristics demonstrated shortened durations for both NREM and REM sleep bouts in knockout mice (Fig. 3d,e,j,k). Concurrently, the duration of wake bouts was prolonged over the 24-hour period (Fig. 3f,l), contributing to the overall disruption of continuous sleep. These results suggest that HSF1 plays a vital role in regulating sleep continuity and stability. *Hsf1* knockout mice exhibited frequent awakenings, indicating difficulty in maintaining sleep states. Collectively, these findings point to HSF1 as a critical regulator of sleep continuity, with its absence leading to fragmented sleep patterns characterized by more frequent but shorter sleep episodes and extended periods of wakefulness.

**Figure 3:**
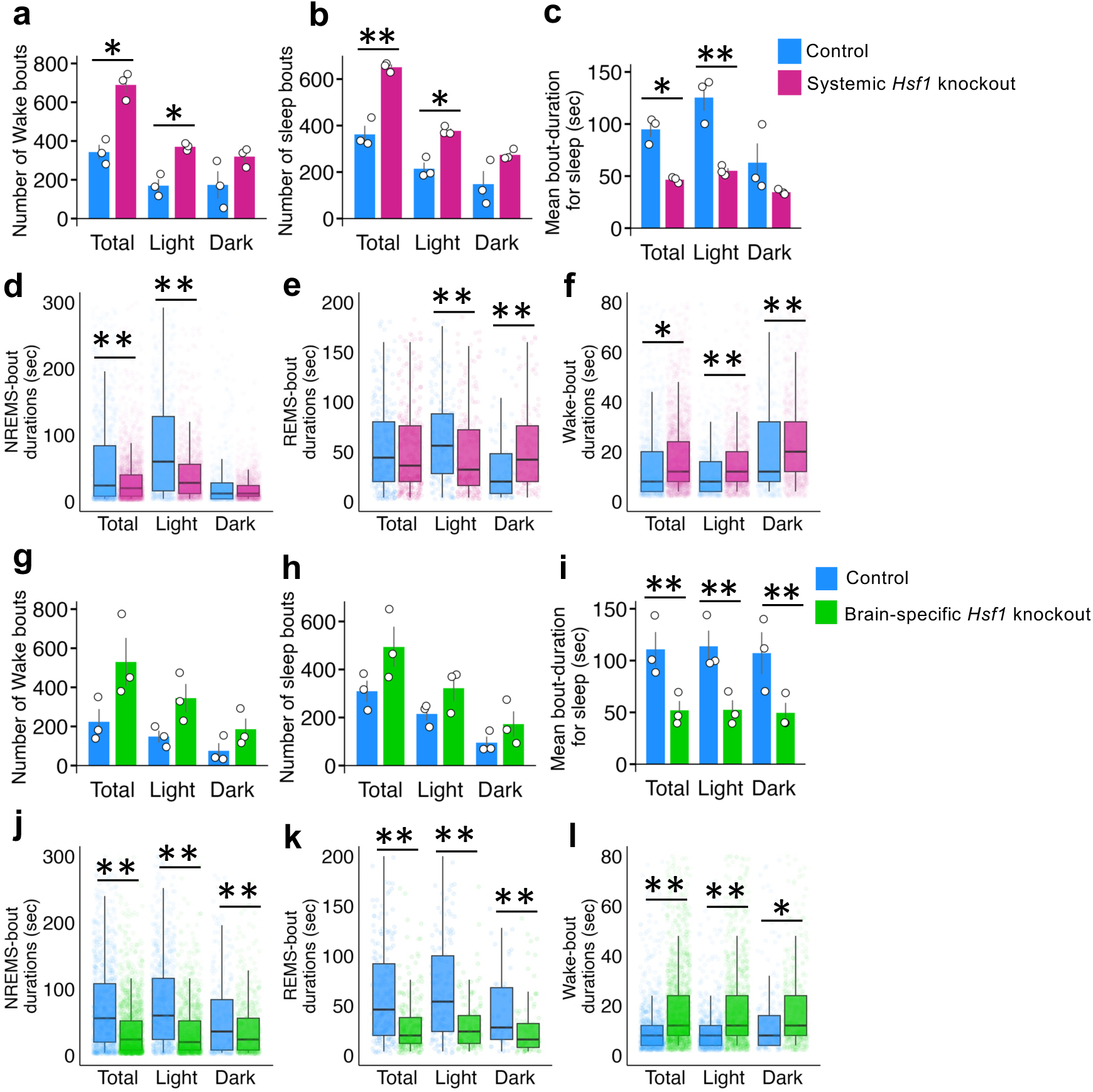
*Hsf1* knockout mice exhibit fragmented sleep. **(a), (g)** Bar plots showing the total number of wake-bouts in each phase (Total: ZT0-24, Light: ZT0-12 or Dark: ZT12-24) for the systemic (a) or brain-specific (g) knockout model. n=3 per group. **(b), (h)** Bar plots showing the mean number of sleep bouts: consecutive (epochs ≥ 2) NREM and REM sleep in each phase for systemic (b) or brain-specific *Hsf1* knockout model (h). **(c), (i)** Bar plots showing the mean bout duration for sleep (seconds): consecutive (epochs ≥ 2) NREM and REM sleep in each phase for systemic (c) or brain-specific (i) *Hsf1* knockout model. **(d), (e), (f), (j), (k), (l)** Box plots showing distributions of bout durations (seconds) for consecutive (epochs ≥ 2) NREM sleep (d,j), REM sleep (e,k), or Wake (f,l) in each phase for systemic (d,e,f) or brain-specific (j,k,l) *Hsf1* knockout model. Data are mean ±s.e.m. with two-way ANOVA with Tukey’s HSD results (a,b,c,g,h,I), median with interquartile range (IQR) and individual data points with Wilcoxon rank-sum test (d,e,f,j,k,l). *: p<0.05, **: p<0.01.

#### Hsf1 knockouts do not show increased NREM sleep after sleep deprivation

Having established the role of *Hsf1* in maintaining sleep continuity, we next investigated how its absence affects the homeostatic response to sleep deprivation (SD) (Fig. 4a). Both systemic and brain-specific *Hsf1* knockout mice showed increased wakefulness and decreased NREM sleep following 6 hours of SD, in contrast to control mice which increased their sleep to compensate for sleep loss (Fig. 4b,c,j,k). Total time spent in wake or NREM sleep during the 6 hours after SD differed significantly between genotypes (Fig. 4d,e,l,m). Although *Hsf1* knockout mice did not show increased NREM sleep, they maintained unfulfilled sleep demands as indicated by consistently higher delta power density in NREM sleep compared to controls (Fig. 4f,n). Both knockout models exhibited a sleep homeostatic response, evidenced by increased delta power during the 2 hours post-SD relative to their baseline (Fig. 4g-i,o-q). However, this was less pronounced in brain-specific knockouts (Fig. 4q).

**Figure 4:**
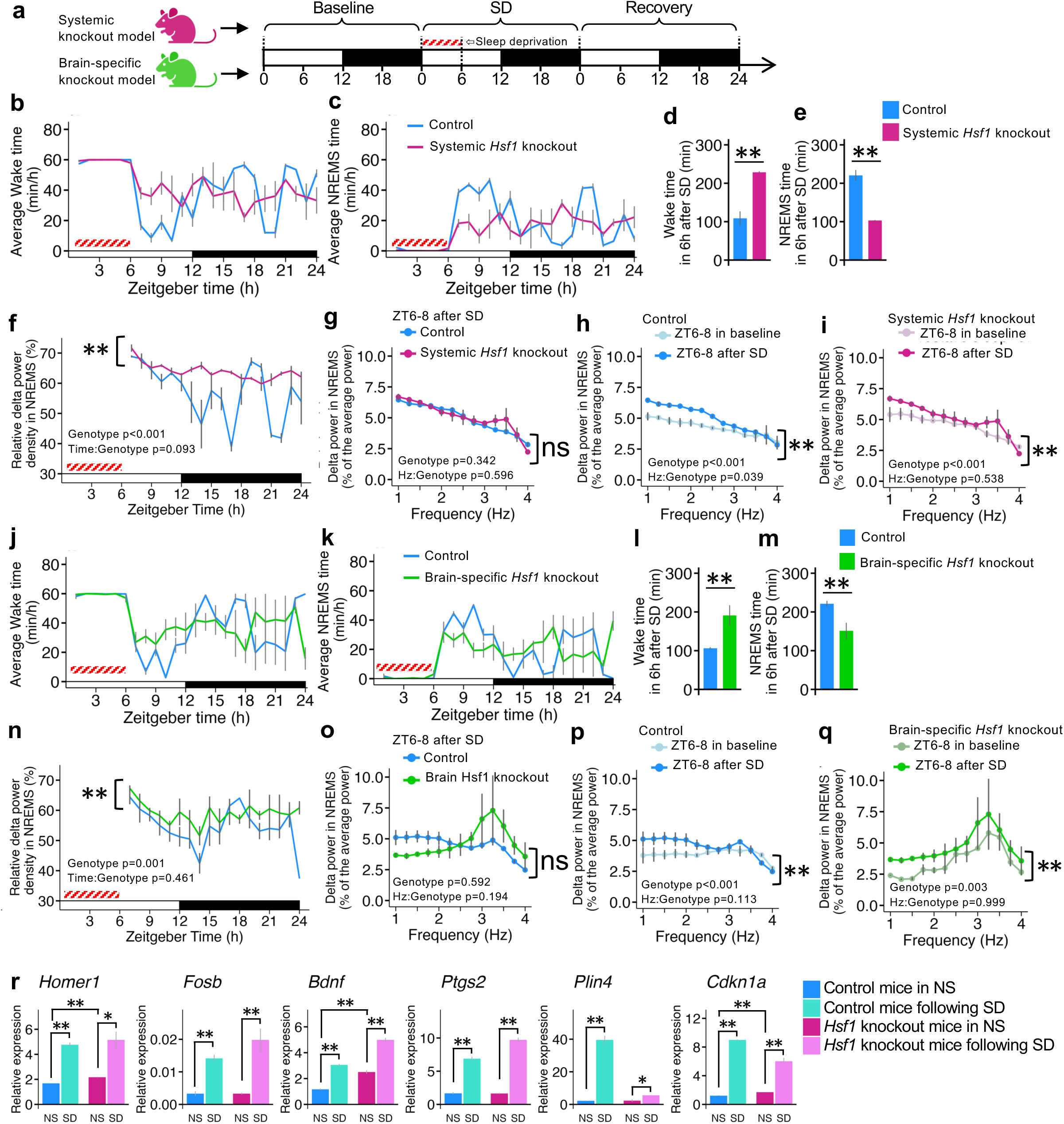
*Hsf1* knockout mice show homeostatic sleep response without increasing NREM sleep following sleep deprivation. **(a)** Schematic of EEG/EMG recording protocol including 6 hours sleep deprivation (ZT6-8 of SD period). **(b), (c), (j), (k)** Line plots showing the hourly average time (minutes) spent in wake (b,j) or NREM sleep (c,k) in SD period for the systemic (b,c) or brain-specific (j,k) knockout model (n=3 per group). **(d), (e), (l), (m)** Bar plots showing the total time (minutes) spent in wake (d,l) or NREM sleep (e,m) in 6 hours after sleep deprivation (ZT6-12 of SD period). **(f), (n)** Line plots showing hourly average of delta power (1-4Hz) density in NREMS (%) in SD period for the systemic (f) or brain-specific (n) knockout model. **(g), (h), (i), (o), (p), (q)** Average delta power density in NREM sleep (%) in 2 hours after sleep deprivation (ZT6-8 of SD period) of controls and knockouts (g,o), baseline and after sleep deprivation of control group (h,p), or of knockouts group (I,q) for the systemic (g,h,i) or brain-specific (o,p,q) knockout model, comparing Control and *Hsf1* knockout group in 1-4Hz range. **(r)** Bar plots showing the relative mRNA expression of *Homer1*, *Fosb*, *Bdnf*, *Ptgs2*, *Plin4* or *Cdkn1a* normalized to *Actb*, in cerebral cortex tissues of *CAG^CreERTM^;Hsf1^flox/flox^*mice treated with corn oil (Control) or tamoxifen (*Hsf1* knockout), harvested at ZT6 during normal sleep (NS) or after 6 hours sleep deprivation (SD) (n=3 per group). Data are mean ±s.e.m. Statistical significance was determined using two-way ANOVA with Tukey’s HSD (f,g,h,I,n,o,p,q), two-tailed t-test (d,e,l,m,r). *: p<0.05, **: p<0.01, ns: not significant. P values for main effect of genotype factor: control or *Hsf1* knockout (Genotype), and interaction effect between genotype and time factors: every hour from ZT1 to ZT24 (Time:Genotype) (f,g,h,i,n,o,p,q).

qPCR analysis of sleep-related genes provided additional evidence of a molecular homeostatic response to SD in both controls and *Hsf1* knockout mice. We examined the expression of a suite of genes that have been reported to be up-regulated in response to sleep deprivation *Homer1, Fosb*, *Bdnf* (*Brain-derived neurotrophic factor*), *Ptgs2* (*Prostaglandin-endoperoxidase synthase 2*), *Plin4* (*Perilipin 4*) and *Cdkn1a* (*Cyclin-dependent kinase inhibitor 1a*) ^24–31^. Expression of these genes increased following SD, suggesting a homeostatic response (Fig. 4r). Interestingly, *Hsf1* knockout mice showed altered expression of some genes (*Homer1*, *Bdnf*, *Cdkn1a*) even under normal sleep conditions, suggesting ongoing sleep disruption (Fig. 4r). Thus, despite the molecular evidence of heightened sleep pressure, *Hsf1* knockout mice were unable to sustain NREM sleep.

The discrepancy between both electrophysiological (delta power) and molecular markers of sleep need and the inability of knockout mice to maintain sleep suggests that HSF1 plays a critical role in sleep regulation beyond simply sensing sleep pressure. Specifically, these findings indicate that HSF1 is essential for the ability to initiate and maintain sleep in response to homeostatic demands, underlining its importance in executing the sleep process itself.

#### Hsf1 deficiency disrupts sleep independently of thermoregulation

We found that *Hsf1* knockout mice showed some differences in their body temperature patterns compared to controls (Supplementary Fig. 4a-l). This raised an important question: are the sleep abnormalities in *Hsf1* knockout mice directly caused by the absence of HSF1, or are they an indirect result of altered temperature regulation? However, the inconsistent and often small temperature differences between knockouts and controls, coupled with similar sleep changes in both systemic and brain-specific knockouts despite their distinct temperature profiles (Supplementary Fig. 4a-c and 4g-i), argue against temperature as the driver of NREM sleep disruptions in *Hsf1* knockouts.

To gain a more fine-grained analysis of temperature-sleep relationships in the knockouts, we conducted a detailed analysis comparing sleep stage transitions with corresponding body temperature changes (Supplementary Fig. 4m,n). We found weak correlations between sleep stage transitions and body temperature changes across all mice. The strongest correlations were during transitions from REM to NREM sleep (Supplementary Fig. 4n). However, these transitions occur infrequently, limiting their significance and broader applicability of this observation. Taken together, these results suggest that the altered sleep patterns in *Hsf1* knockout mice are a direct consequence of HSF1’s role in sleep regulation, rather than an indirect effect of disrupted temperature regulation ^32,33^.

#### HSF1 enhances synaptic organization to maintain sleep

To elucidate the molecular mechanisms by which HSF1 maintains normal sleep patterns, we performed CUT&RUN ^34^ and bulk RNA sequencing (RNA-seq) on cerebral cortex from C57BL/6J mice under normal sleep (NS) and sleep deprivation (SD) conditions (Fig. 5a). RNA-seq revealed numerous differentially expressed genes (DEGs) between NS and SD conditions (Fig. 5b), while CUT&RUN identified genes directly regulated by HSF1 genomic binding (Fig. 5c). By comparing these datasets, we identified 1421 DEGs from RNA-seq, 1528 HSF1-regulated genes from CUT&RUN, and 178 genes that overlapped between the two methods (Fig. 5d). This overlap represents genes directly regulated by HSF1 in response to sleep deprivation.

**Figure 5:**
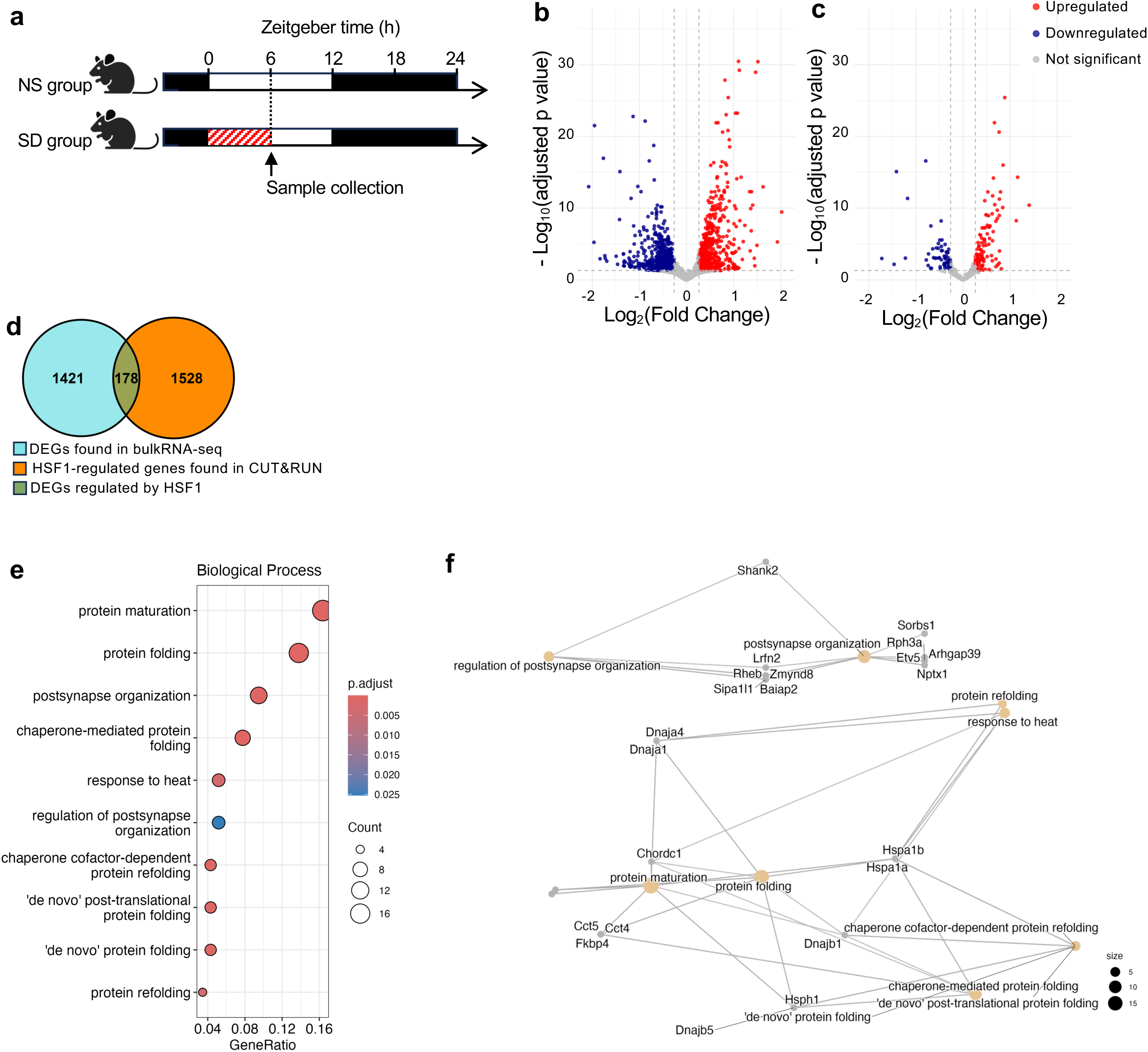
HSF1 enhances synaptic organization during sleep deprivation. **(a)** Schematic showing the protocol of sleep deprivation and sample collection used for transcriptomic analysis. Red diagonal strip pattern indicates the sleep deprivation (ZT0-6). **(b)** Volcano plot showing the genes found in bulk-RNA seq using mRNA from cerebral cortex tissues of normal sleep (NS) or sleep deprived (SD) group (n=3 mice per group). The vertical axis indicates -log_10_(P value) from Wilcoxon rank-sum test with Bonferroni correction. The horizontal axis indicates log_2_(fold change) values. **(c)** Volcano plot showing the genes found both in CUT&RUN and bulkRNA-seq. (cutoff: adjusted p-value<0.05 and 1.2<|fold change|). **(d)** Venn diagram showing the total number of differentially expressed genes (DEGs) during SD which identified by bulkRNA-seq analysis (1421 genes) and genes associated with the HSF1 binding loci increased during SD which identified by CUT&RUN analysis (1528 genes). Overlapping area indicates the DEGs regulated by HSF1 (178 genes). **(e)** Enriched biological processes by Gene Ontology (GO) analysis on the up-regulated genes by HSF1. (cutoff: adjusted p-value<0.05 and q<0.05). **(f)** Network plot (Cnetplot) illustrating the relationships between enriched biological processes and genes that are significantly upregulated in SD group. The nodes represent either genes or biological processes, with gene nodes connected to the biological process nodes if they are associated with a particular GO term. The size of the biological process nodes correlates with the number of genes involved in each process.

Gene Ontology (GO) analysis of HSF1-regulated DEGs revealed significant enrichment of genes related to protein folding and chaperone functions, as expected. Importantly, it also showed enriched biological processes in postsynapse organization and regulation of postsynapse organization (Fig. 5e). Gene network analysis further illustrated connections between HSF1-regulated genes involved in synaptic organization (Fig. 5f). These findings suggest that HSF1 plays a vital role in orchestrating transcriptional programs that maintain synaptic functions, in addition to its known role in the heat shock response. This mechanism appears to be essential for sustaining sleep, as evidenced by the inability of *Hsf1* knockout mice to maintain sleep states.

#### Hsf mutation in flies disrupts continuous sleep despite increased demand

HSF1 is highly conserved across species. This conservation suggests a fundamental role in cellular processes, potentially including sleep regulation ^8,35^. To investigate its evolutionary role in sleep, we extended our studies to *Drosophila* melanogaster, which has a single ortholog, *Hsf*. This diurnal model organism is well-established in sleep research ^36,37^. We compared sleep patterns between *Hsf* mutant flies and matched genetic controls under three conditions: baseline, sleep deprivation (SD), and recovery (Fig. 6a).

**Figure 6:**
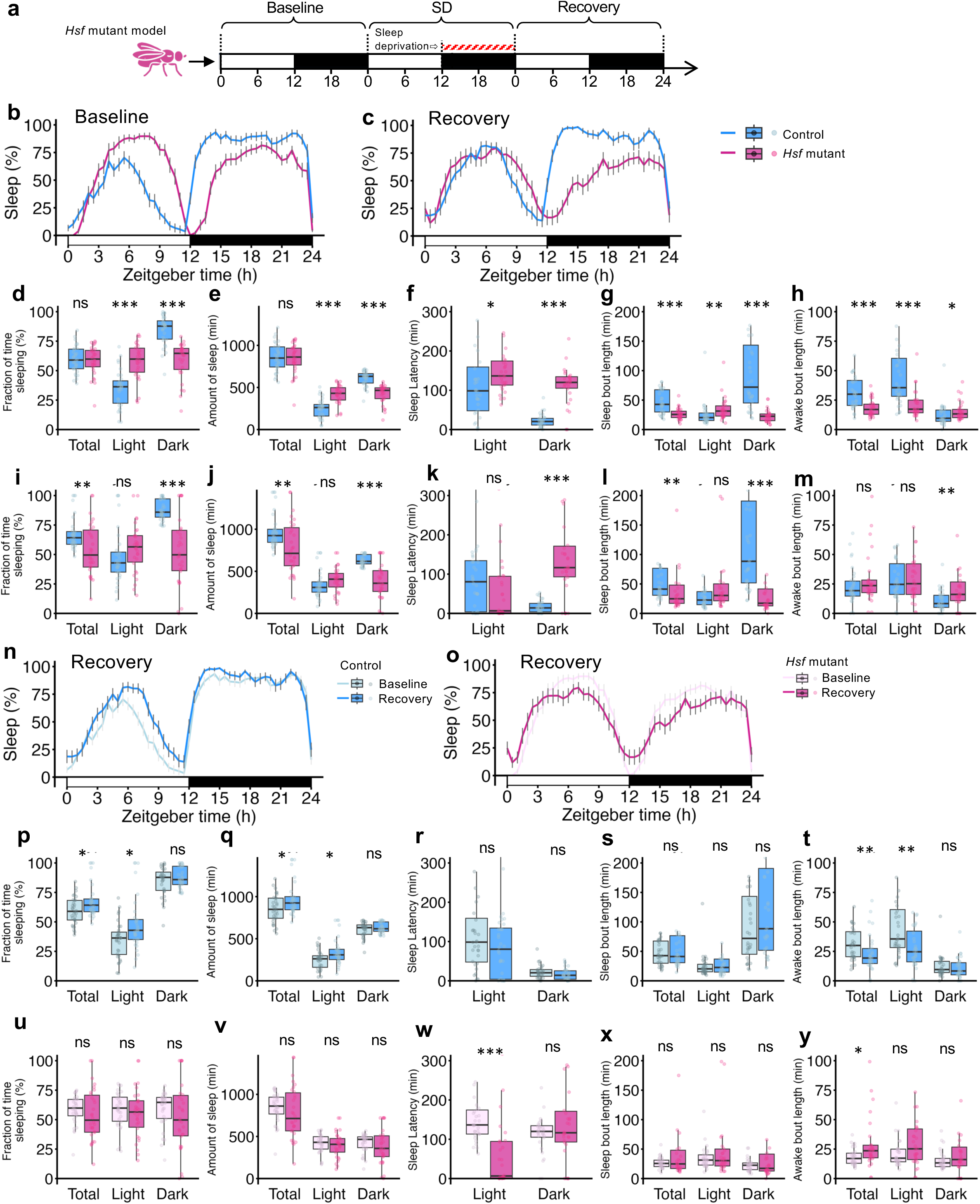
*Hsf* mutation in flies disrupts continuous sleep despite increased sleep demand. **(a)** Schematic of locomotor activity recording protocol using *Drosophila melanogaster*. Locomotor data were collected in 1 minute bin, and inactivity for at least 5 minutes was defined as a sleep bout. Dead flies were excluded from the analysis (control group: n= 32, *Hsf* mutant group: n= 30). Results for Baseline (b,d,e,f,g,h) or Recovery (c,i,j,k,l,m) period comparing control and mutant group (b,c,d,e,f,g,h,I,j,k,l,m), or Baseline and Recovery periods in control (n,p,q,r,s,t) or mutant group (o,u,v,w,x,y). **(b), (c), (n), (o)** Line plots showing the flies’ average proportion of time spent sleeping in consecutive 30 minutes segments (%) over the 24-hour period in 12h-light/12h-dark cycle. **(d), (i), (p), (u)** Box plots showing distributions of the average proportion of time spent sleeping in consecutive 30 minutes segments (%) during Total (ZT0-24), Light (ZT0-12) or Dark (ZT12-24) phase. (**e), (j), (q), (v)** Box plots showing distributions of the mean sleep amount (minutes) during each phase. **(f), (k), (r), (w)** Box plots showing distributions of the mean latency (minutes) to the first sleep event from ZT0 (Light) or ZT12 (Dark). **(g), (l), (s), (x)** Box plots showing distributions of the average bout-length (minutes) for sleep during each phase. **(h), (m), (t), (y)** Box plots showing distributions of the average bout-length (minutes) for awake during each phase. Data are mean ±s.e.m. (b,c,n,o) or median with IQR and data points for individual average (d,e,f,g,h,I,j,k,l,m,p,q,r,s,t,u,v,w,x,y). Statistical significance was determined using two-step approach (d,e,f,g,h,I,j,k,l,m,p,q,r,s,t,u,v,w,x,y): First, data for each group (control or *Hsf* mutant) and phase (Total, Light or Dark) were assessed for normality using the Shapiro-Wilk test. If the data were normally distributed (p ≥ 0.05), an independent samples t-test was employed to compare the means between independent groups within each phase. If the data significantly deviated from normality (p < 0.05), the Mann-Whitney U test, a non-parametric alternative, was used. All tests were conducted separately for different phases and conditions. *: p<0.05, **: p<0.01, ***: p<0.001, ns: not significant.

Under baseline conditions, *Hsf* mutant flies maintained typical daily sleep patterns. They exhibited daytime napping and nighttime sleep (Fig. 6b). However, mutants showed an increased daytime sleep fraction and reduced nighttime sleep fraction. Total 24-hour sleep remained unchanged (Fig. 6d,e). This suggests a compensatory shift in sleep distribution due to disrupted regulation. Mutant flies struggled with sleep initiation and maintenance. They showed extended latency to the first nighttime sleep bout and reduced nighttime sleep bout length (Fig. 6f,g). Daytime awake bouts were shorter in mutants (Fig. 6h). This may reflect an increased need for daytime naps to compensate for insufficient nighttime sleep.

To assess how *Hsf* mutation affects sleep recovery mechanisms, we next subjected flies to 12h sleep deprivation during their normal sleep phase (ZT12-24; night). In the first 12 hours post-deprivation, both mutant and control flies showed no significant differences in sleep behavior (Fig. 6c,i-m). However, comparing baseline and recovery sleep within each genotype revealed distinct characteristics of *Hsf* mutant flies (Fig. 6n,o). Control flies exhibited an increased sleep fraction and reduced awake bout length during daytime recovery (Fig. 6p,q,s,t). In contrast, mutant flies showed no increase in sleep length (Fig. 6u,v,x,y). Mutants did, however, exhibit increased sleep demand after deprivation, evidenced by reduced latency to the first sleep bout during the light phase (Fig. 6r,w).

Collectively, these findings parallel our observations in *Hsf1* knockout mice. *Hsf* mutant flies are compromised in their ability to increase sleep amount despite elevated sleep demand. The disturbance in continuous sleep and difficulties converting sleep demand to sleep execution highlight a conserved role of HSF1. Thus, this transcription factor appears essential for maintaining sleep stability across species.

## Discussion

Our results reveal a critical role for HSF1 in sleep regulation, extending beyond its well-established function in cellular stress responses ^38,39^. We demonstrate that HSF1 is dynamically regulated in the brain during sleep-wake cycles and is essential for maintaining sleep continuity and quality. Oscillation of nuclear HSF1 levels in the mouse brain, with increases during wakefulness and decreases during sleep, suggests a direct involvement of HSF1 in sleep-wake regulation. This aligns with previous studies showing dynamic gene expression changes during sleep-wake cycles and highlights the importance of transcriptional regulators in sleep homeostasis ^40,41^. The upregulation of nuclear HSF1 during sleep deprivation further supports its role in responding to extended wakefulness.

Our use of both systemic and brain-specific *Hsf1* knockout mouse models provides strong evidence for HSF1’s direct involvement in sleep regulation. Both models exhibited significant reductions in NREM sleep and increased wakefulness, particularly during the light phase. This phenotype suggests that HSF1 is vital for initiating and maintaining sleep states. The fragmented sleep patterns observed in *Hsf1* knockouts, characterized by more frequent but shorter sleep episodes, further emphasize HSF1’s role in sustaining continuous sleep. These findings are consistent with reports that disruptions in synaptic integrity, which are suggested by our molecular studies in *Hsf1* knockouts, may lead to fragmented sleep ^17,42–44^. Intriguingly, *Hsf1* knockout mice maintained normal sleep pressure, as evidenced by persistent delta power and upregulation of sleep-related genes like *Homer1 and Bdnf* after sleep deprivation. However, *Hsf1*-deficient mice struggled to increase NREM sleep in response to sleep loss. This discrepancy between sleep need and sleep execution suggests that HSF1 is essential for translating sleep pressure into actual sleep. It remains unclear whether it may also contribute to the regulation of sleep need.

Our molecular analyses provide insight into the mechanisms by which HSF1 regulates sleep. CUT&RUN and RNA-seq data reveal that HSF1 regulates genes involved in synaptic organization. This finding also expands our understanding of HSF1’s functions beyond traditional heat shock responses and suggests an alternative mechanism by which it maintains sleep stability. The enrichment of HSF1-regulated genes in postsynaptic organization pathways implies that HSF1 may orchestrate transcriptional programs essential for synaptic function and plasticity during sleep.

HSF1’s role in sleep regulation is conserved across species. Our findings in *Drosophila* mirror those in mice, with *Hsf* mutant flies showing disrupted sleep patterns. These include fragmented nighttime sleep and increased daytime sleep. This cross-species consistency suggests evolutionarily ancient molecular mechanisms. Previous studies support this idea. Transcriptional regulation of synaptic proteins is crucial for sleep homeostasis in *Drosophila* ^45,46^. This points to a shared mechanism of HSF1’s effects between mice and flies. These findings open new avenues for understanding the complex interplay between HSF1, synaptic regulation, and sleep continuity across species.

## Conclusions

Our findings reveal HSF1 as a critical regulator of sleep, primarily through its influence on synaptic organization. This study not only expands our understanding of HSF1’s biological functions but also provides new insights into the molecular underpinnings of sleep regulation. The identification of HSF1 as a key player in sleep maintenance opens new avenues for investigating sleep disorders and potentially developing targeted therapeutic interventions to improve sleep quality and overall health.

## Materials and Methods

### Mice

All animal studies were carried out in concordance with an approved protocol from the Institutional Animal Care and Use Committee (IACUC) at Perelman School of Medicine at the University of Pennsylvania. Male C57BL/6J mice were purchased from Jackson Laboratory. All the mice used were at the age of 6-8 weeks when the experiments began. Before sleep deprivation, mice were singly housed for habituation with *ad libitum* access to food and water under standard humidity and temperature (21 ± 1 °C) on a 12-h light: 12-h dark (dim red light < 10 lux) cycle.

### Construction of conditional Hsf1 knockout mouse

To generate a conditional knockout model of *Hsf1*, we utilized CRISPR/Cas9-mediated genome editing combined with single guide RNAs (sgRNAs) targeting exon 2 of the *Hsf1* gene, in conjunction with a floxed single-stranded DNA (ssDNA) template for homology-directed repair ^47^. We designed two sgRNAs to target exon 2 of the *Hsf1* gene: Mm_Hsf1_Ex2_Left:5’- ACACCAUCAUAGUUUCACUG-3’ and Mm_Hsf1_Ex2_Right:5’-CAUCUCUUAGAAAUAGGCUG-3’ (CRISPRevolution sgRNA EZ Kit、Synthego). Each sgRNA was resuspended by dissolving in injection buffer (IDTE pH 7.5, 11-01-02-02, Integrated DNA Technologies Inc.), and stored at - 80°C until use. The ssDNA template was designed to introduce LoxP sites flanking exon 2 of *Hsf1*. This was resuspended in nuclease-free water and stored at -80°C until use. For the preparation of the CRISPR/Cas9 microinjection mix, the sgRNAs were combined with Alt-R^TM^ S.p. HiFi Cas9 Nuclease V3 (1081060, Integrated DNA Technologies Inc.) to form the ribonucleoprotein (RNP) complexes. We mixed each of sgRNAs with Cas9 and injection buffer to the final concentration of each sgRNA 10 ng/µL, and the Cas9 concentration 50 ng/µL in the injection mix. This mixture was incubated at room temperature for 10 minutes to allow for the formation of RNP complexes. To the RNP complex mix, the ssDNA was added to achieve a final ssDNA concentration of 10 ng/µL per kilobase. The mix was filtered through Ultrafree Centrifugal Filter (UFC30VV25, MilliporeSigma) to remove particulates, and the filtrate was collected as the final injection mix. The CRISPR/Cas9 RNP and ssDNA injection mix was kept on wet ice until microinjection. Standard microinjection procedures were followed to inject the mixture into the pronuclei of C57BL/6J zygotes (RRID:IMSR_JAX:000664). Injected zygotes were transferred into pseudopregnant recipient females to establish founder mice harboring the floxed *Hsf1* allele. Genotyping of the founder (F0) mice was performed by PCR to confirm the successful insertion of LoxP sites flanking exon 2 of the *Hsf1* gene: *Hsf1^flox/flox^*. Genotyping primers were as follows: Primer_1: 5’- CACTGCTCAGACCTAGGTTCCATGG-3’, Primer_2: 5’-GGTCAAACACGTGGAAGCTGTTCC-3’, Primer_3: 5’-ACAACAACATGGCTAGCTTCGTGC-3’, Primer_4: 5’- AGCTTCCCTTTTTGTAAATGGAAGTGTGG-3’, Primer_5: 5’-GAGAGACATCAGGCCACAGATAACTTCG-3’, Primer_6: 5’-CTCCATCTCTTAGAAATAGGATAACTTCGTATAATGTATGC-3’. To produce the experimental *Hsf1* knockout line, we followed these breeding steps. Generation of F1: An inbred cross of F0 *Hsf1^flox/flox^*founders was performed to produce F1 *Hsf1^flox/flox^* offspring. Generation of F2: F1 *Hsf1^flox/flox^* mice were backcrossed with pure background C57BL/6J mice (RRID:IMSR_JAX:000664), yielding F2 *Hsf1^flox/+^*heterozygous mice. Generation of F3: An inbred cross of F2 *Hsf1^flox/+^*mice was conducted to obtain F3 *Hsf1^flox/flox^* offspring. Generation of F4: F3 *Hsf1^flox/flox^* mice were crossed with *CAG^CreERTM^*transgenic mice which have a tamoxifen-inducible Cre-mediated recombination system driven by the chicken beta actin promoter/ enhancer coupled with the cytomegalovirus (CMV) immediate-early enhancer: B6.Cg-Tg(CAG-cre/Esr1*)5Amc/J (RRID:IMSR_JAX:004682), resulting in F4 *Hsf1^flox/+^;CAG^CreERTM^*mice. Generation of F5: F4 *Hsf1^flox/+^;CAG^CreERTM^* male were crossed with *Hsf1^flox/flox^* females to produce F5 *Hsf1^flox/flox^;CAG^CreERTM^* mice. Expansion for Experiments (F6): F5 *Hsf1^flox/flox^;CAG^CreERTM^* mice were crossed with *Hsf1^flox/flox^* mice, generating F6 *Hsf1^flox/flox^;CAG^CreERTM^* mice for experimental use. For systemic knockouts, *Hsf1^flox/flox^:CAG^CreERTM^* mice were treated with tamoxifen to induce ubiquitous Cre-mediated recombination, which deletes exon 2 in *Hsf1*. Controls were littermates with the same genotype treated with corn oil. For brain-specific knockouts, *Hsf1^flox/flox^*mice were injected with AAV-CAG-Cre-GFP (PHP.eB) virus, leading to CNS-specific deletion of exon 2 of *Hsf1*. Controls were littermates with the same genotype injected with AAV-CAG-GFP (PHP.eB). The conditional *Hsf1* knockout mouse model was developed with support from the Penn Transgenic and Chimeric Mouse Facility.

### Gene modification in mice

Tamoxifen (T5648, Sigma-Aldrich) was dissolved in corn oil (C8267-500ML, Sigma) at 50°C for a final concentration at 20mg/mL and filtered through sterile syringe filter (Minisart 0.2µm PES, S6534-FM6UK, Sartorius Stedim Biotech). *Hsf1^flox/flox^:CAG^CreERTM^* transgenic mice were intraperitoneally injected with the tamoxifen at 150 mg/kg/day for five consecutive days. Mice were used for experiments one week after the final injection. For gene modification using AAVs, mice were anesthetized using isoflurane and injected 5x10^11^ vector genomes (vg) in 100µL of PBS per mouse into the right retro-orbital sinus using 29-gauze Insulin Syringes (1484132, Fisher Scientific). Mice were used for subsequent experiments six weeks after the AAV injection. Verification of *Hsf1* knockout was performed by extracting DNA from cerebral cortex tissues after each treatment, followed by PCR analysis to confirm the excision of the floxed exon 2 in *Hsf1*, and immunoblot and histology analysis to verify a decrease of HSF1 protein in the tissue. The primers used for the PCR, Primer_1: 5’-CACTGCTCAGACCTAGGTTCCATGG-3’, Primer_4: 5’- AGCTTCCCTTTTTGTAAATGGAAGTGTGG-3’, Primer_5: 5’-GAGAGACATCAGGCCACAGATAACTTCG-3’, Primer_6: 5’- CTCCATCTCTTAGAAATAGGATAACTTCGTATAATGTATGC-3’, were obtained from Sigma-Aldrich.

### Sleep phenotyping

To monitor brain and muscle activity over an extended period, we employed a telemetric approach using HD-X02 transmitters (Data Sciences International) implanted in mice between 6 and 11 weeks of age. The surgical procedure was conducted under isoflurane anesthesia (3–4% for induction, 2–2.5% for maintenance). We implanted two EEG electrodes (2.4 mm shaft length, 2.16 mm head diameter, 1.19 mm shaft diameter; Plastics One) epidurally above the right frontal and parietal cortices, following established protocols ^13,48^. These electrodes were linked to the transmitter using medical-grade stainless steel wires and protected with dental cement (867-1601, Reliance). For EMG recording, we inserted two stainless-steel leads into the neck musculature, spacing them about 5 mm apart and securing them with sutures. The transmitter itself was positioned subcutaneously along the left dorsal flank. To manage post-operative discomfort, we administered buprenorphine (Vetergesic, 0.1 mg/kg) and meloxicam (Metacam, 10 mg/kg) subcutaneously at the start of surgery. Following a recovery period of at least 10 days, we initiated experimental protocols. Continuous EEG/EMG data collection was performed over 6–7 days using specialized hardware and Dataquest ART software (Data Sciences International). The neurophysiological signals were transmitted at 455 kHz to an RPC-1 receiver and digitized at a 250 Hz sampling rate.

### Sleep Scoring and Spectral Analysis

Vigilance states were categorized for consecutive 4-second epochs through visual examination of EEG and EMG signals using SleepSign for Animal ver. 3 (Kissei Comtec) as detailed previously ^13^. We defined wakefulness as periods with high and variable EMG activity accompanied by low-amplitude EEG signals. Non-rapid eye movement sleep (NREMS) was characterized by high-amplitude EEG dominated by slow waves and low-amplitude EMG. Rapid eye movement sleep (REMS) was identified by low EEG amplitude, theta oscillations in the 5–9 Hz range, and absence of EMG muscle tone. We computed EEG power spectra for each 4-second epoch using Welch’s method in Python, analyzing frequencies from 0.25 to 25 Hz with 0.25 Hz resolution and applying a Hanning window function. Epochs containing EEG artifacts were excluded from spectral analyses, accounting for 7.9 ± 0.8% of total recording time. EEG power spectra were calculated separately for NREMS, REMS, and wakefulness during the 12-hour light and dark periods across 3 recording days. We expressed these spectra as percentages of total EEG power (1–25 Hz range, 1 Hz resolution). EEG delta power was computed by summing power in the 1–4 Hz range, both in total and during NREMS specifically. This metric was averaged over consecutive intervals containing equal numbers of 4-second NREM sleep epochs and then normalized as a percentage of levels observed between ZT8-12 on day 1 (baseline).

### Sleep Deprivation

For sleep recording (EEG/EMG), baseline recording, 24 hours before sleep deprivation, was used as a control condition after 1-week of habituation period in each cage. Sleep deprivation was initiated as lights on (ZT0) the next day and performed for 6 hours by gentle handling using a brush to prevent sleep ^29,49,50^. After sleep deprivation (ZT6), mice were kept in the same cage undisturbed until ZT24, which were used as SD sleep. Recording on the following day (ZT0-ZT24) was classed as recovery sleep. For brain sample collection, 6 hours sleep deprivation (ZT0-ZT6) was performed by a gentle handling after 2 weeks of acclimatization. The samples were flash-frozen in liquid nitrogen and stored at -80°C until use.

### Drosophila lines

In this study, we utilized two *Drosophila melanogaster* lines: the *Hsf* mutant (cn[1] bw[1] Hsf[4], DGRC_108256) expressing a mutant HSF protein (V57M) with reduced DNA-binding activity ^51,52^, obtained from the Drosophila Genetic Resource Center, and its genetic background control (cn[1] bw[1], BDSC_264) from the Bloomington Drosophila Stock Center. Flies were maintained on a standard food medium (0.7% agar, 2.0% inactivated yeast, 5.0% molasses; Fly Food J, 7013-PNV, LABExpress) at 18°C with a 12h-light:12h-dark cycle. For experiments, we selected virgin females, initially using 32 flies per group. Locomotor activities were recorded using DAM2 Drosophila Activity Monitor (TriKinetics Inc.), with individual flies placed in glass tubes containing food. Their movement was monitored in 12h-light:12h-dark cycles at 25°C, with data collected at 1-minute intervals and analyzed for activity patterns, sleep duration, and circadian rhythms. Flies were acclimated for 24 hours before data collection. Following baseline recording, flies underwent sleep deprivation using a custom-programmed mechanical stimulus during the 12-hour dark phase, consisting of 2-second pulses every 20 seconds with a 10% random sigma applied to avoid habituation. After 12 hours of sleep deprivation, a 24-hour recovery period was implemented without stimuli.

### Fly sleep analyses

Sleep was defined as periods of inactivity lasting five or more minutes. Activity data were processed using damr (R package version 0.3.7), behavr (R package version 0.3.2), ggetho (R package version 0.3.7), scopr (R package version 0.3.3), sleepr (R package version 0.3.0), zeitgebr (R package version 0.3.5) ^53^ in R Statistical Software (version 4.3.1; R Foundation for Statistical Computing). Dead flies were excluded if their activity counts were less than 10 counts during the recovery period. The data were converted into 30-minute intervals for the analysis of average sleep fraction, calculated for each genotype and replicate. The analysis was separated into light (ZT0-12) and dark (ZT12-24) phases to assess sleep behaviors under different environmental conditions. The percentage of time spent sleeping was calculated for each 30- minute interval over the total (ZT0-24), light, and dark phases. The average sleep percentages for each interval were plotted to visualize sleep dynamics over time. Statistical comparisons between control and *Hsf* mutant flies were made for both baseline, SD and recovery phases. For clarity, the data were summarized as mean percentages with SEM for each phase, and sleep fraction box plots were generated. Sleep and awake bout lengths were computed for baseline, SD, and recovery phases. Sleep bouts were defined as periods of inactivity lasting at least five minutes. Mean, median, and standard deviation of bout lengths were calculated across all phases. Sleep latency was determined by calculating the time from lights on (ZT0) or lights off (ZT12) to the first sleep bout, defined as more than five consecutive sleep bouts, in each phase.

### Cell lines

HEK293T cells were cultured in Dulbecco’s Modified Eagle’s Medium (DMEM; D6030, Sigma-Aldrich) containing 10% Fetal Bovine Serum (F2442, Lot 21B456, Sigma-Aldrich) and 1% Penicillin Streptomycin (15140122, Gibco) in an incubator with a humidified atmosphere at 37°C containing 5% CO_2_.

### Adeno-associated virus production

HEK293T cells were seeded in DMEM supplemented with 10% FBS and 1% Penicillin Streptomycin the day before the transfection. 60% confluent HEK293T cells were triple transfected with pAdDeltaF6, pUC-mini-iCAP-PHP.eB and expression plasmid at the same molarity using polyethyleneimine (43896, Thermo Scientific Chemicals). 90% of the culture medium were replaced by OptiPRO (12309019, Fisher Scientific) supplemented with 1% Penicillin Streptomycin at 24 hours after the transfection. The cells were cultured in the medium for additional 72 hours. Viruses were extracted using chloroform and precipitated in 50% PEG (Polyethylene glycol). AAVs were concentrated using AmiconUltra4 Centrifugal Filter Unit (UFC810024, Millipore) and stored at 4°C until use. The virus titer was measured by quantitative PCR on serial diluted samples using PowerUp SYBR Green Master Mix (Applied Biosystems) and primers against the ITRs, forward: 5’-GGAACCCCTAGTGATGGAGTT-3’, reverse: 5’-CGGCCTCAGTGAGCGA-3’, obtained from Sigma-Aldrich.

### Immunohistochemistry (IHC)

Fresh frozen brain tissue was sectioned using a cryostat at 10µm thickness and fixed by immersing slides in acetone for 7 minutes at -20°C, followed by methanol for 7 minutes at -20°C, then allowing the agents to evaporate at room temperature for 30 minutes. The sections were blocked in TBST containing 5% Donkey serum (D9663, Sigma) for 1 hour at room temperature. Primary antibodies, including Anti-Neun Antibody, clone A60 (MAB377, Abcam), Polyclonal Anti-Glial Fibril Acid Protein (Z033429-2, Abcam), and anti-HSF1 antibody (4356, Cell Signaling), were diluted in blocking buffer (500, 1000, and 500 times, respectively) and applied to the slides overnight at 4°C. Secondary antibodies, including Goat anti-Mouse IgG(H+L) Cross-Absorbed Secondary Antibody, Alexa Fluor 568 (A-11031, Invitrogen), Donkey anti-Rabbit IgG(H+L) Cross-Absorbed Secondary Antibody, Alexa Fluor 488 (A-21206, Invitrogen), and Donkey anti-Rabbit IgG(H+L) Cross-Absorbed Secondary Antibody, Alexa 647 (A-31573, Invitrogen), were applied and incubated for 1 hour at room temperature. The samples were then mounted using ProLong Glass Antifade Mountant with NeucBlue (P36981, Invitrogen) and Cover glass (631-0137, VWR). Fluorescent signals were detected using an ECLIPSE Ti microscope and NIS Elements AR 4.13.04 software (Nikon).

### Immunoblotting

Whole brains harvested from mice were flash-frozen and stored at -80°C until use. Nuclear extractions were performed using the Minute Single Nucleus Isolation Kit for Tissue/Cells (SN-047, Invent Biotechnologies, Inc.) following the manufacturer’s instructions. Total cell and extracted nuclei were lysed by incubation in modified RIPA buffer (50mM Tris-HCl (pH7.4), 150mM NaCl, 1% Triton X-100, 1% Sodium deoxycholate, 0.25% SDS, with protease and phosphatase inhibitors) for 10 minutes on ice, followed by sonication (10 cycles, 30 seconds on/1 minute off). Protein samples were then reduced using Pierce DTT (20291, Thermo Scientific) and heated in Laemmli Sample Buffer (161-737, BioRad) at 95°C for 10 minutes. 50µg of protein per lane were loaded on NuPAGE Novex 4-12% Bis-Tris gradient gels (WG1403BOX, Invitrogen) and run in 1X Bolt MES SDS Running Buffer (B000202, Invitrogen) at 200V for 35 minutes. Proteins were transferred to nitrocellulose membranes (IB301031, Invitrogen) using iBlot (Invitrogen). After a brief wash, membranes were blocked for 1 hour in 0.5% BSA/nonfat dried milk in 1X TBST (J77500-K2, Sigma-Aldrich) and incubated with Monoclonal Anti-U2AF65 antibody (U4758, MilliporeSigma) at 1:2000 dilution in blocking buffer overnight at 4°C. Membranes were washed three times for 10 minutes each in TBST and incubated with secondary antibodies (Anti-Mouse IgG-Peroxidase; A4416, Sigma-Aldrich, or Goat anti-Rabbit IgG-HRP; A16110, Invitrogen) at 1:10,000 dilution for 2 hours at room temperature. After three 10-minute washes, membranes were incubated with Immunoblot for 5 minutes. Chemiluminescent signals were detected using an ImageQuant LAS 4000 and Control Software Version 1.3 (GE Healthcare).

*RNA extraction and quantitative reverse transcription polymerase chain reaction (qRT-PCR)* Total RNA was extracted from fresh frozen mouse cerebral cortex tissues using Direst-zol RNA miniprep (R2052, Zymo Research). The extracted RNA was reverse-transcribed into cDNA using the High-Capacity cDNA Reverse Transcription Kit (4368814, Applied Biosystems) following the manufacturer’s instructions. qRT-PCR was performed using TaqMan Gene Expression Master Mix (4369016, Applied Biosystems) and commercially available primers for m1 Hsf1 (Mm01201402), m1 Per2 (Mm00478113), m1 ARNTL1 (Mm00500226), Homer1 (Mm00516275_m1), Fosb (Mm00500401_m1), Bdnf (Mm04230607_s1), Ptgs2 (Mn00478374_m1), Plin4 (Mm00491061_m1), Cdkn1a (Mm04205640_g1), and s1 Actb (Mm00607939). The RT-PCR reactions were run in duplicate for each target gene using a ViiA7 Real-Time PCR System (Applied Biosystems). Expression levels of target genes were determined using the delta-delta cycle threshold method to calculate fold changes, with normalization to beta actin expression levels.

#### CUT&RUN

Harvested cerebral cortex from mice were flash-frozen and store at -80°C until use. Intact nuclei were extracted used for CUT&RUN. Minced cerebral cortex tissues were incubated in the Nuclear Extraction Buffer: 20mM HEPES (15630080, Gibco), 10mM Potassium chloride (P9541, Sigma-Aldrich), 20% (v/v) Glycerol (G5516, Sigma-Aldrich), 0.1% NP-40 (85124, Thermo Scientific), 0.5mM Spermidine (S0266, Sigma-Aldrich), 1X Halt Protease Inhibitor Cocktail (78425, Fisher Scientific) and 2% Bovine Serum Albumin (B6917, Sigma-Aldrich), for 10 minutes on ice and filtered through 40µm CellPro Premium Cell Strainer (CS4040, DOT SCIENTIFIC INC.). These steps were repeated three times before proceeding the following workflows. CUT&RUN wes performed according to the manufacture’s instruction. In brief, activated Concanavalin A Conjugated Paramagnetic Beads (211401, ConA beads, EpiCypher) were incubated with the extracted nuclei to bind for 10 minutes at room temperature. The ConA beads-nuclei mixture was resuspended in cold Antibody Buffer: 20mM HEPES (Gibco), 150mM Sodium chloride (S5150, Sigma-Aldrich), 0.5mM Spermidine (Sigma-Aldrich), 1X Halt Protease Inhibitor Cocktail (Fisher Scientific), 0.05% Digitonin (300410, Millipore Sigma), 2mM EDTA (15575020, Invitrogen) and 2% Bovine Serum Albumin (Sigma-Aldrich), and incubated with 0.5µg of anti-HSF1 antibody (4356S, Cell Signaling), SNAP-ChIP Certified H3K4me3 Positive Control Antibody (MA511198, Thermo Fisher) or Rabbit IgG Negative Control Antibody (130042, EpiCypher) per 5x10^5^ nuclei at 4°C overnight on nutator. After washing twice in cold Digitonin Buffer: 20mM HEPES (Gibco), 150mM Sodium chloride (S5150, Sigma-Aldrich), 0.5mM Spermidine (Sigma-Aldrich), 1X Halt Protease Inhibitor Cocktail (Fisher Scientific), 0.05% Digitonin (300410, Millipore Sigma) and 2% Bovine Serum Albumin (Sigma-Aldrich), the ConA beads-nuclei-antibody mixtures were resuspended in Digitonin Buffer and incubated with 2.5µL of CUTANA pAG-MNase (151116, EpiCypher) per 5x10^5^ nuclei for 1 hour at 4°C on nutator for binding. The mixtures were washed twice, resuspended in Digitonin Buffer with 2mM Calcium chloride (21115, Sigma-Aldrich) and incubated at 4°C for 2 hours for chromatin digestion. After the incubation, Stop Buffer: 340mM Sodium chloride (Sigma-Aldrich), 20mM EDTA (Invitrogen), 4mM EGTA (E8145, Sigma-Aldrich), 50µg/mL RNase A (EN0531, Thermo Scientific), 50µg/mL Glycogen (10901393001, Roche), 0.5mM Spermidine (Sigma-Aldrich), 1X Halt Protease Inhibitor Cocktail (Fisher Scientific), 0.02% Digitonin (300410, Millipore Sigma) and 2% Bovine Serum Albumin (Sigma-Aldrich), was added and incubated for 10 minutes at 37°C to stop the digestion. The cleaved chromatins were collected from the supernatant and purified using Monarch PCR&DNA Cleanup Kit (T1030L, BioLabs) according to the manufacture’s instruction.

### CUT&RUN Library preparation and sequencing

Sequencing libraries for each condition were prepared using 10ng of CUT&RUN enriched DNA fragments in total made by pooling equal amount DNAs of three replicates. End repair was performed using NEBNext Ultra II End Repair/dA-Tailing Module (E7546L, New England BioLabs) according to the manufacturer’s instructions. Then, the samples were ligated to the NEBNext Adaptors (E6440L, New England BioLabs) and cleaned up using 1.75X (v/v) SPRIselect beads (B23318, Beckman Coulter). Adaptor-ligated DNA fragments were amplified using NEBNext Unique Dual Index Primer Pairs (New England BioLabs, E6441A) with NEBNext Ultra II Q5 Master Mix (M0544S , New England BioLabs). PCR was performed using the following parameters: initial denaturation at 98°C for 30 seconds, 14 cycles of denaturation at 98°C for 10 seconds and annealing/extension at 65°C for 75 seconds, final extension at 65°C for 5 minutes. Size selections for 150-500bp were performed on the PCR products by applying 0.6X and 1.2X SPRIselect beads successively. Sequencing using a NovaSeq X Plus 10B flow cell platform with paired-end 300- 500bp reads and subsequent quality filtering of reads was performed according to the manufacturer’s instructions (Illumina).

### CUT&RUN bioinformatics analysis

FASTQ files containing sequence reads were processed with CUT&RUNTools2.0. Peak calling for ant-HSF1 antibody was performed using control, corresponding aligned data for IgG negative control antibody in each replicate, with MACS2 (version 2.2.9.1) using standard threshold setting (q<0.05). The peak calling results were annotated with transcription starting sites (TSS; within ± 3,000bp rage from gene) using ChIPseeker (R package version 3.38.0) along with BSgenome.Mmusculus.UCSCmm39 (R package version 1.4.3), and the TSS-associated genes were listed in each replicate. Genes associated with HSF1-binding loci increased during sleep deprivation (increased-genes in SD group) were determined by subtracting the genes found in all the replicates of NS group from the genes found in one or more replicates in SD group.

### bulkRNA sequencing (bulkRNA-seq) sample and library preparation

Total RNA was extracted using Direct-zol RNA miniprep (R2052, Zymo Research) from fresh frozen mouse cerebral cortex tissues. messenger RNA (mRNA) was isolated using NEBNext Poly(A) Magnetic Isolation Module (E7490L, New England BioLabs) according to the manufacture’s instruction. RNA-seq libraries were prepared using NEBNext Ultra II RNA Library Prep Kit for Illumina (E7770, New England BioLabs). End repair was performed using NEBNext Ultra II End Repair/dA-Tailing Module (E7546L, New England BioLabs) according to the manufacturer’s instructions. Then, the samples were ligated to the NEBNext Adaptors (E6440L, New England BioLabs) and cleaned up using 1.8X (v/v) SPRIselect beads (B23318, Beckman Coulter). Adaptor-ligated DNA fragments were amplified using NEBNext Unique Dual Index Primer Pairs (E6441A, New England BioLabs) with NEBNext Ultra II Q5 Master Mix (M0544S, New England BioLabs). PCR was performed using the following parameters: initial denaturation at 98°C for 30 seconds, 7 cycles of denaturation at 98°C for 10 seconds and annealing/extension at 65°C for 75 seconds, final extension at 65°C for 5 minutes. Size selections for 150-500bp were performed on the PCR products by applying 0.9X and 1.2X SPRIselect beads successively. The libraries were quantified by qPCR using NEBNext Library Quant Kit for Illumina (E7642S, New England BioLabs) according to the manufacture’s instruction, and mixed in equal weight proportion to form the sequence library. Sequencing using a NovaSeq X Plus 10B flow cell platform with paired-end 300-500bp reads and subsequent quality filtering of reads was performed according to the manufacturer’s instructions (Illumina).

### bulkRNA-seq bioinformatics analysis

FASTQ files were trimmed using TrimGalore (version 0.5.0) with parameters: clip first 40 bases of both forward and reverse readas, then aligned to the STAR index built on GRCm39 using STAR (version 2.7.11b). Counts, FPKM and TPM were computed using Stringtie (version 2.2.1) with options: -e -B -G. The read count information was extracted and compiled to a table by running prepDE.py3 script with default setting. Using the computed gene count matrix above, following analysis was performed on R Statistical Software. Expression count of at least 1 across samples were retained for analysis. DESeq2 (R package version 1.44.0) analysis was applied to the count object comparing NS and SD group. Shrunken log_2_ fold changes (LFC) and SE were calculated using lfcShrink with the apeglm method on the DESeq2 results ^54^. Then the HSF1-regulated genes were determined as the overlapping genes found both in the DESeq2 results and the increased- genes in SD group from CUT&RUN analysis. Those genes were filtered with the cutoff values: adjusted p value<0.05 and |fold change|<1.2. Gene ontology analysis was performed on the filtered genes using enrichGO function from clusterProfiler ^55^ (R package version 3.0.4) with the cutoff values: adjusted p value<0.05 and q value<0.05. The results were plotted using dotplot and cnetplot functions from enrichplot (R package version 1.24.0).

### Statistical analyses

The statistical significance of differences between two groups was assessed using a two-step approach: normality was first evaluated with the Shapiro-Wilk test. If data were normally distributed (p≥0.05), a two-tailed t-test was used: otherwise, the Mann-Whitney U test was applied. One-way analysis of variance (ANOVA) was applied to comparisons with multiple data points. Two-way ANOVA with Tukey’s HSD post-hoc test was applied to multiple comparisons with two or more independent factors. Spearman’s rank correlation coefficient was applied to detect the strength and direction on a relationship between two variables. Results were presented with significance considered at *P* < 0.05. Statistical software R was used to perform these analyses.

## Acknowledgements

A.B.R. acknowledges funding from the Perelman School of Medicine, University of Pennsylvania, the Institute for Translational Medicine and Therapeutics (ITMAT) at the University of Pennsylvania. This work was supported also by NIH DP1DK126167 and R01GM139211 (A.B.R.).

## Author contributions

S.Y. performed animal experiments, manual sleep stage scoring, performed the molecular and histological analysis techniques, produced virus, prepared the sequencing samples and libraries, analyzed results and wrote the manuscript. U.K.V. performed fly experiments. analyzed experimental results and designed and implemented model code. J.C-R. performed mouse breeding and fly experiments. A.B.R. designed the project and analyzed results, secured funding, and wrote the paper with contributions from the other authors.

---

**Supplementary Figure 1:**
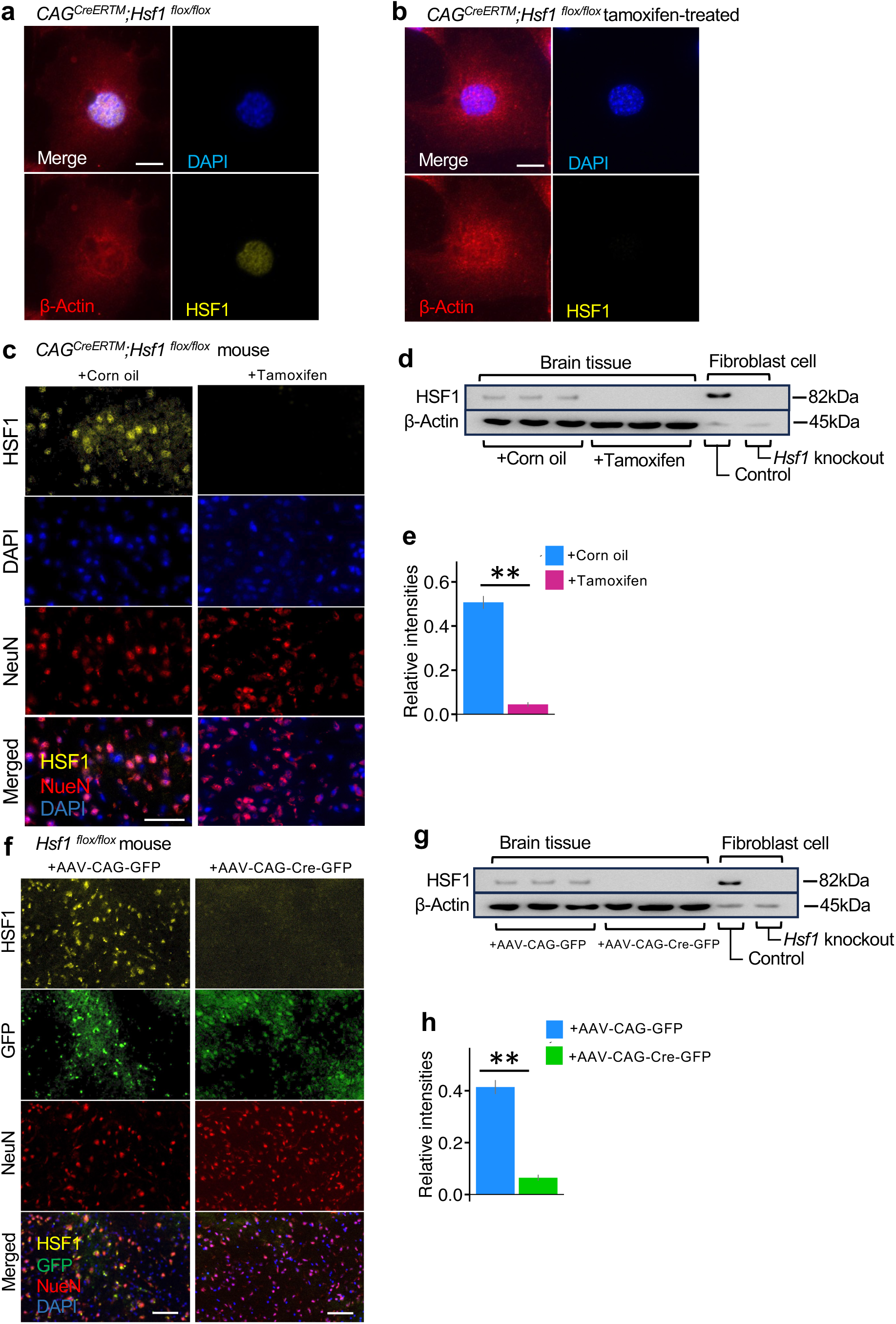
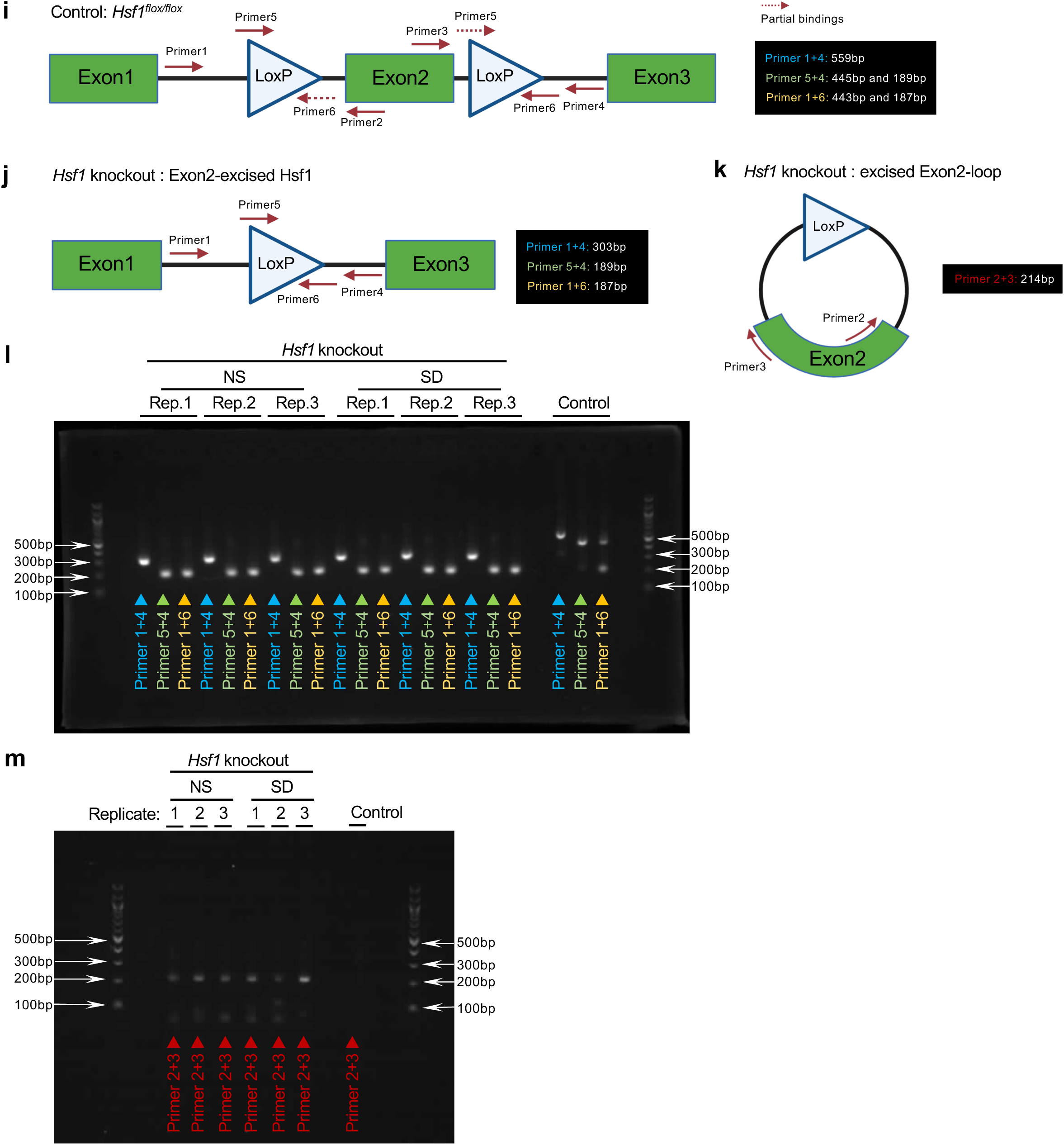
Validation of *Hsf1* knockout. **(a)**, **(b)** Immunohistochemistry showing HSF1 expression in skin fibroblast cells from a mouse (*CAG^CreERTM^;Hsf1^flox/flox^*) without treatment (a), or with tamoxifen-treatment (b). Scale bars: 25µm. (**c), (f)** Immunohistochemistry showing HSF1 expressions in the cerebral cortex of systemic *Hsf1* knockout model: *CAG^CreERTM^;Hsf1^flox/flox^*(c) or brain-specific *Hsf1* knockout model: *Hsf1^flox/flox^*(f). Scale bars: 200µm. **(d), (g)** Immunoblotting showing HSF1 expressions in the cerebral cortex of systemic *Hsf1* knockout model (d) or brain-specific *Hsf1* knockout model (g). **(e), (h)** Bar plots showing the relative signal intensities of HSF1 expression normalized to beta-actin expression in the brain tissues of systemic *Hsf1* knockout model (e) or brain-specific *Hsf1* knockout model (h). **(i), (j), (k)** Schematics showing the structure of *Hsf1^flox/flox^*region (i), *Hsf1^flox/flox^* region after exon 2 excision at the LoxP sites by Cre-recombinase activity (j), extra-chromosomal circular DNA containing the excised exon 2 (k), and the primer binding sites used for genotyping. Expected PCR product sizes are listed for each primer combination on each of the gene structure. **(l)** Gel image showing the truncation of exon 2 in the *Hsf1* knockout mice: PCR products of DNA extracted from cerebral cortex tissues using the primer combinations: Primer1 and 4, Primer5 and 4, or Primer1 and 6. **(m)** Gel image showing the existence of extra-chromosomal circular DNA containing exon 2: PCR products of DNA extracted from cerebral cortex tissues using Primer2 and 3. Data are mean ±s.e.m. Statistical significance was determined using two-tailed t-test (e,h). **: p<0.01.

**Supplementary Figure 2:**
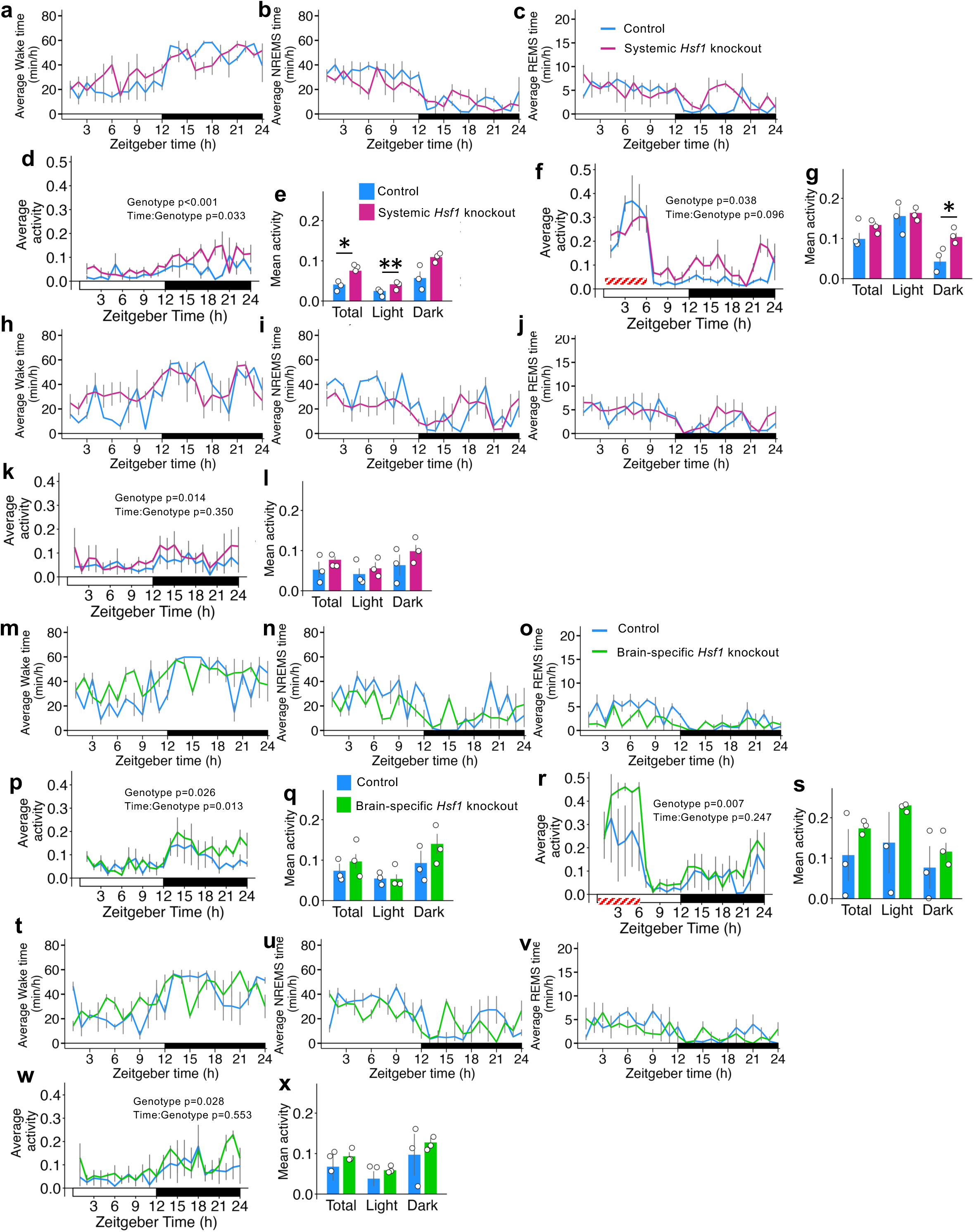
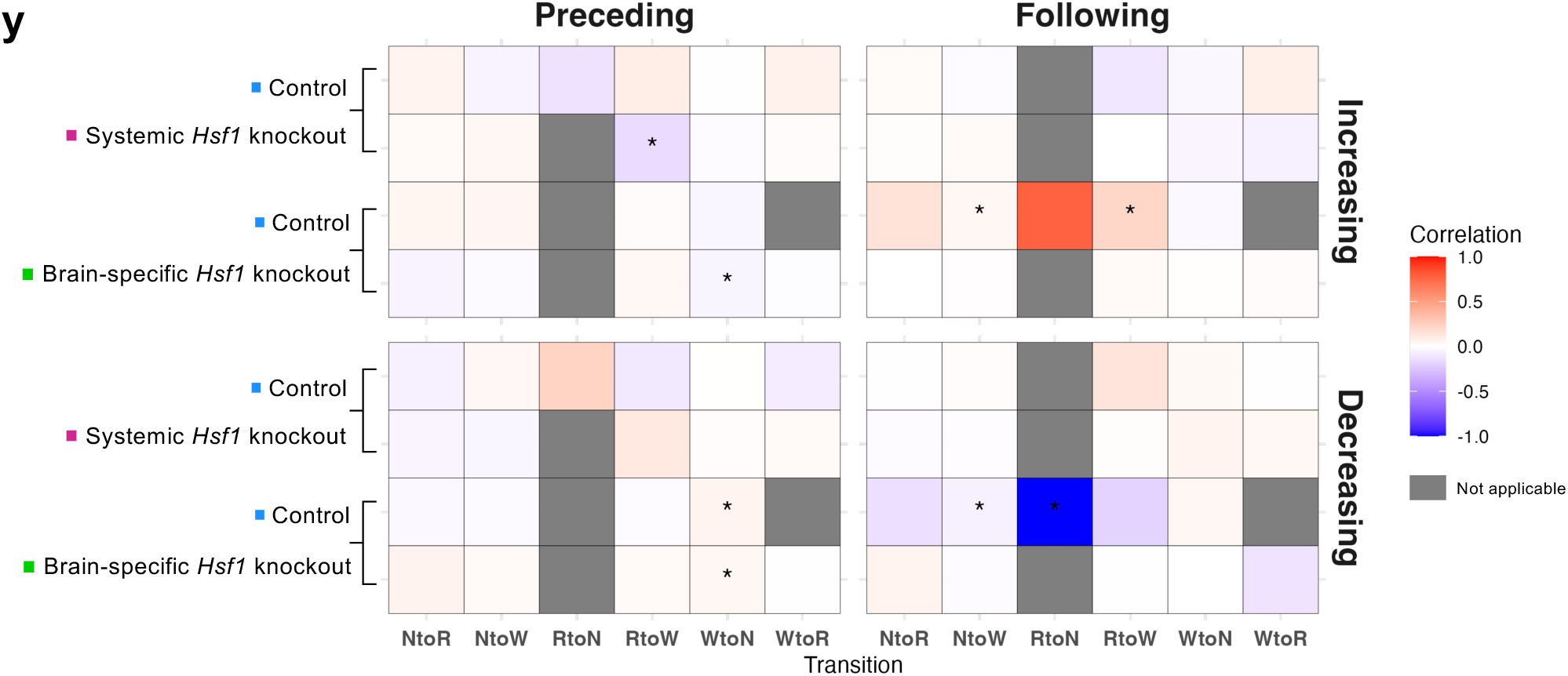
Sleep architectures and locomotor activities of *Hsf1* knockout mice. **(a), (b), (c), (h), (i), (j), (m), (n), (o), (t), (u), (v)** Line plots showing hourly average time (minutes) spent in wake (a,h,m,t), NREM sleep (b,i,n,u) or REM sleep (c,j,o,v) during Baseline (a,b,c,m,n,o) or Recovery (h,i,j,t,u,v) period for systemic *Hsf1* knockout models (a,b,c,h,i,j) or brain-specific *Hsf1* knockout models (m,n,o,t,u,v). n=3 per group. **(d), (f), (k), (p), (r), (w)** Line plots showing hourly average of activity counts (counts per 100 milliseconds) in Baseline (d,p), SD (f,r) or Recovery (k,w) period for the systemic *Hs1* knockout model (d,f,k) or brain-specific *Hsf1* knockout model (p,r,w). **(e), (g), (l), (q), (s), (x)** Bar plots showing mean activity counts (counts per 100 milliseconds) during Total (ZT0-24), Light (ZT0-12) or Dark (ZT12-24) phase in Baseline (e,q), SD (g,s) or Recovery (l,x) period for the systemic *Hs1* knockout model (e,g,l) or brain-specific *Hsf1* knockout model (q,s,x). **(y)** Heat maps visualizing the correlation between activity change (Increasing or Decreasing) and occurrence timing (Preceding or Following) around various sleep stage transitions during Baseline period in four groups: *CAG^CreERTM^;Hsf1^flox/flox^*mice treated with corn oil (Control). *CAG^CreERTM^;Hsf1^flox/flox^*mice treated with tamoxifen (Systemic *Hsf1* knockout). *Hsf1^flox/flox^* mice treated with AAV-CAG-GFP (Control). *Hsf1^flox/flox^* mice treated with AAV-CAG-Cre-GFP (Brain-specific *Hsf1* knockout). X-axis represents different sleep stage transitions, including NREM sleep to Wake (NtoW), NREM sleep to REM sleep (NtoR), REM sleep to Wake (RtoW), Wake to NREM sleep (WtoN), and Wake to REM sleep (WtoR). The colors represent the correlation coefficients ranging from -1.0 (blue) to +1.0 (red), not applicable (dark grey). Data are mean ±s.e.m. Statistical significance was determined using two-way ANOVA with Tukey’s HSD (d,e,f,g,k,l,p,q,r,s,w,x), or Spearman’s rank correlation coefficient (y). *: p value<0.05, **: p value<0.01.

**Supplementary Figure 3:**
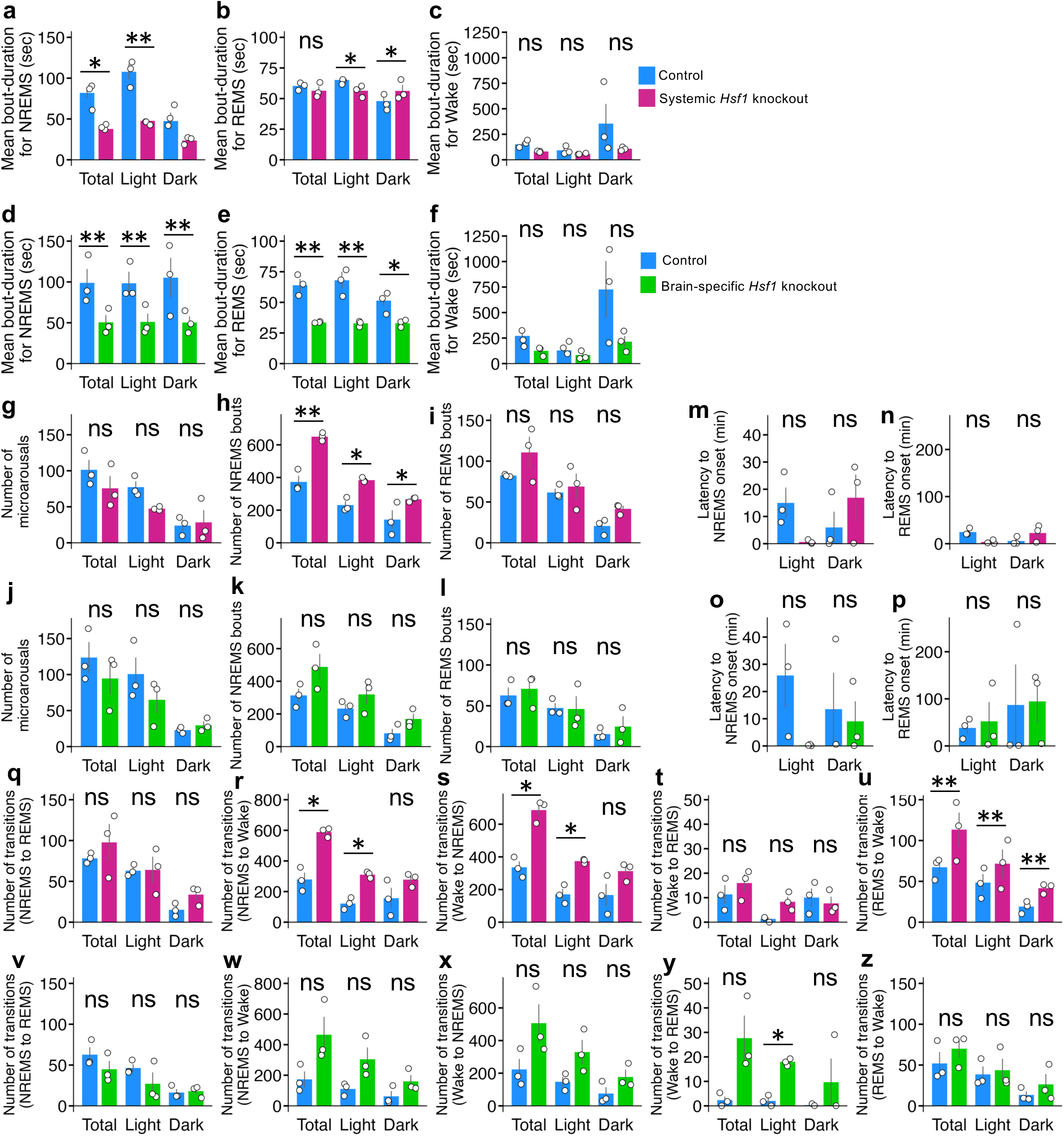
Spectral analysis of *Hsf1* knockout mice during Baseline period. **(a), (b), (c), (d), (e), (f)** Bar plots showing the mean duration (seconds) of bouts (2 or more consecutive epochs) for NREM sleep (a,d), REM sleep (b,e) or Wake (c,f) in each phase (Total: ZT0-24, Light: ZT0-12 or Dark: ZT12-24) during Baseline period for systemic *Hsf1* knockout models (a,b,c) or brain-specific *Hsf1* knockout models (d,e,f). n=3 per group. **(g), (h), (i), (j), (k), (l)** Bar plots showing the average number of bouts for microarousals (g,j), NREM sleep (h,k), REM sleep (i,l) in each phase for systemic *Hsf1* knockout models (g,h,i) or brain-specific *Hsf1* knockout models (j,k,l). **(m), (n), (o), (p)** Bar plots showing the mean latency (minutes) to the first consecutive NREM sleep (m,o) or REM sleep (n,p) in each phase. Sleep latency was defined as the time from ZT0 (start of Light phase), or ZT12 (start of Dark phase) to the detection of first sleep event for systemic *Hsf1* knockout models (m,n) or brain-specific *Hsf1* knockout models (o,p). **(q), (r), (s), (t), (u), (v), (w), (x), (y), (z)** Bar plots showing the average number of sleep-stage transitions in each phase. Transitions from NREM sleep to REM sleep (q,v), NREM sleep to Wake (r,w), Wake to NREM sleep (s,x), Wake to REM sleep (t,y), REM sleep to Wake (u,z) in each phase for systemic *Hsf1* knockout models (q,r,s,t,u) or brain-specific *Hsf1* knockout models (v,w,x,y,z). Data are mean ±s.e.m. Statistical significance was determined using two-way ANOVA with Tukey’s HSD. *: p<0.05, **: p<0.01, ns: not significant.

**Supplementary Figure 4:**
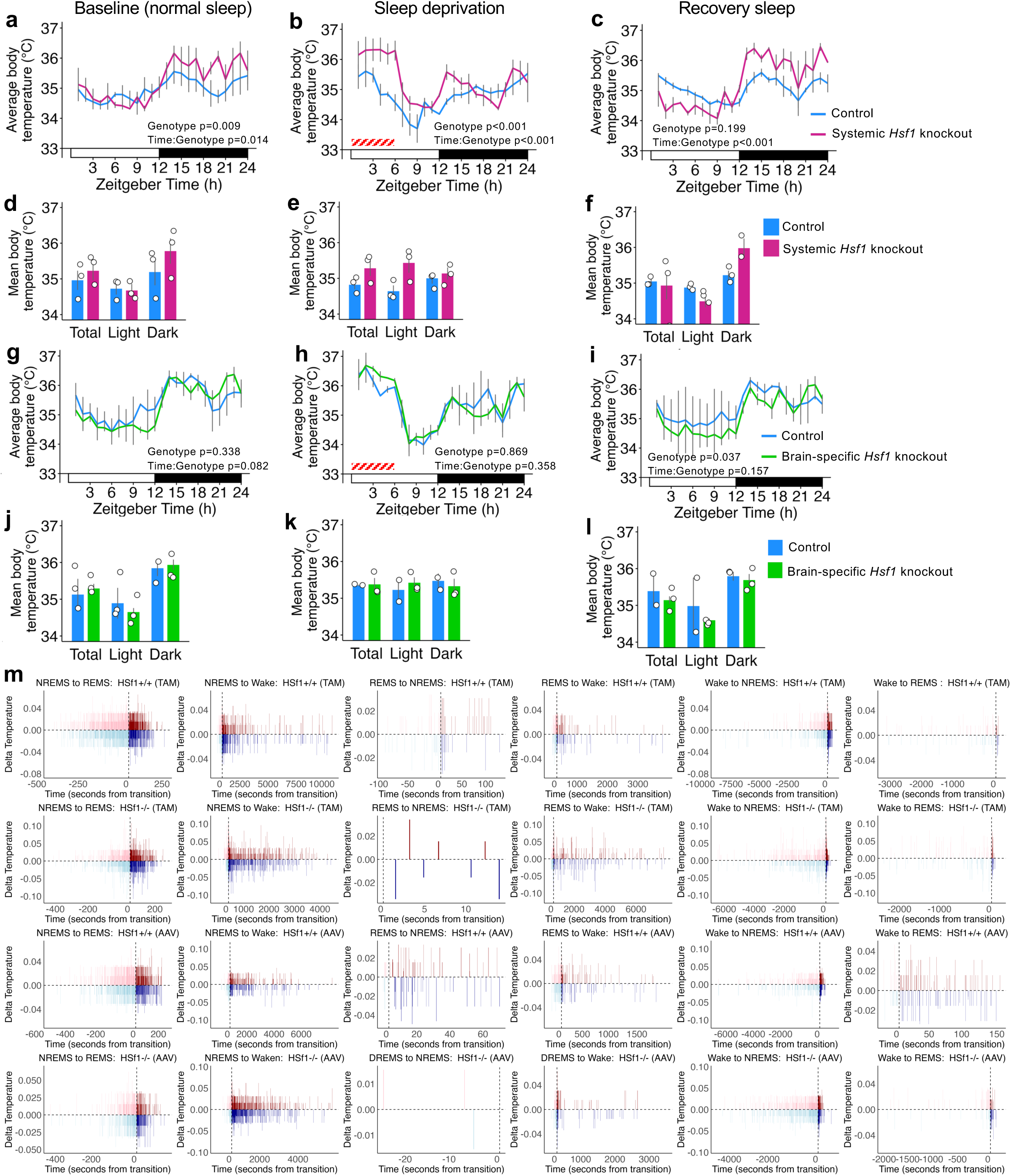
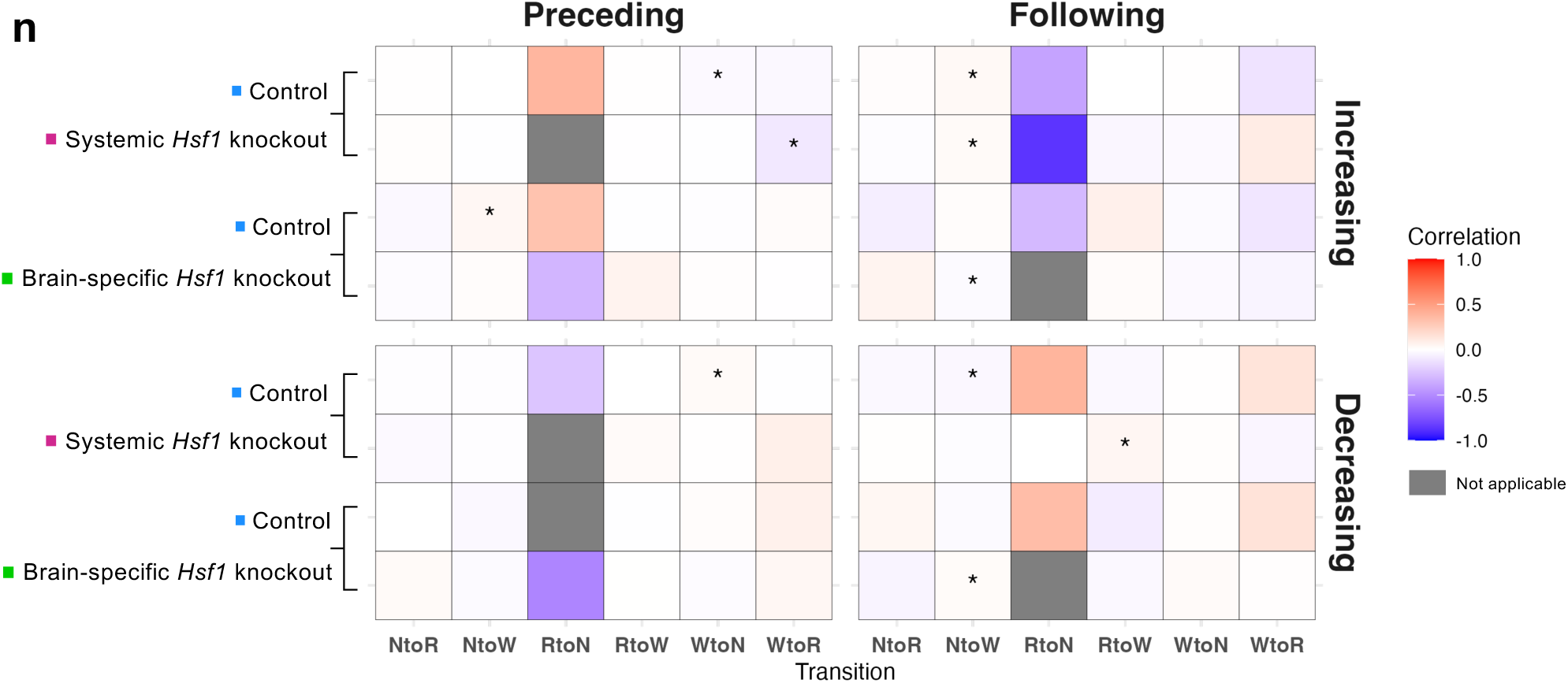
Body temperature and sleep stage transitions of *Hsf1* knockout mice. **(a), (b), (c), (g), (h), (i)** Line plots showing hourly average of body temperature (°C) in Baseline (a,g), SD (b,h) or Recovery (c,i) period for the systemic *Hs1* knockout model (a,b,c) or brain-specific *Hsf1* knockout model (g,h,i), with the p values for main effect of genotype factor: control or *Hsf1* knockout (Genotype) and interaction effect between genotype and time factors: every hour from ZT1 to ZT24 (Time:Genotype). **(d), (e), (f), (j), (k), (l)** Bar plots showing mean body temperatures (°C) during Total (ZT0-24), Light (ZT0-12) or Dark (ZT12-24) phase in Baseline (d,j), SD (e,k) or Recovery (f,l) period for the systemic *Hs1* knockout model (d,e,f) or brain-specific *Hsf1* knockout model (j,k,l). **(m)** Bar plots showing body temperature changes (delta temperature, °C) during the Baseline period occurring around various sleep stage transitions in four groups: Hsf1+/+ (TAM): *CAG^CreERTM^;Hsf1^flox/flox^* mice treated with corn oil (Control). Hsf1-/- (TAM): *CAG^CreERTM^;Hsf1^flox/flox^*mice treated with tamoxifen (Systemic *Hsf1* knockout). Hsf1+/+ (AAV): *Hsf1^flox/flox^* mice treated with AAV-CAG-GFP (Control). Hsf1-/- (AAV): *Hsf1^flox/flox^* mice treated with AAV-CAG-Cre-GFP (Brain-specific *Hsf1* knockout). The time in seconds relative to the transition point is on the x-axis, with the delta temperature changes on the y-axis, where positive values indicate an increase in temperature and negative values indicate a decrease. Bar colors reflect the direction and timing of these temperature changes: dark red for temperature increases after the transition, pink for increases before the transition, dark blue for decreases after the transition, and light blue for decreases before the transition. Only delta temperatures occurring between any preceding and following sleep stage transitions are plotted. **(n)** Heat maps visualizing the correlation between delta temperature (Increasing or Decreasing) and occurrence timing (Preceding or Following) around various sleep stage transitions. X-axis represents different sleep stage transitions, including NREM sleep to Wake (NtoW), NREM sleep to REM sleep (NtoR), REM sleep to Wake (RtoW), Wake to NREM sleep (WtoN), and Wake to REM sleep (WtoR). The colors represent the correlation coefficients ranging from -1.0 (blue) to +1.0 (red), not applicable (grey). Data are mean ±s.e.m. Statistical significance was determined using two-way ANOVA with Tukey’s HSD (a-l), or Spearman’s rank correlation coefficient (n). *: p value<0.05, **: p value<0.01.

**Supplementary Figure 5:**
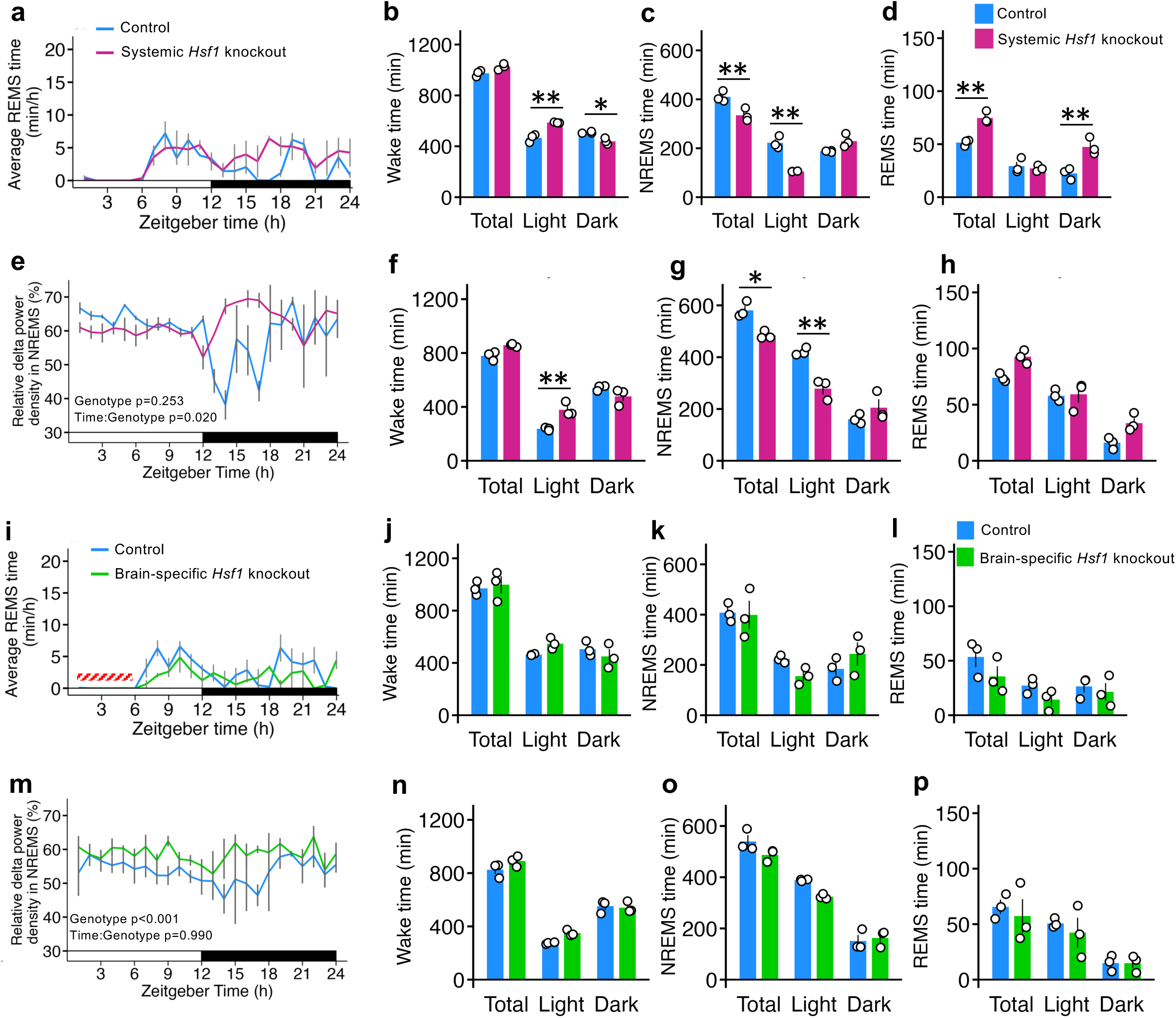
Sleep spectral analysis of systemic and brain specific *Hsf1* knockout mice during SD and Recovery periods. **(a)**, **(i)** Line plots showing the hourly average time (minutes) spent in REM sleep during SD period (n=3 per group) for the systemic *Hs1* knockout model (a) or brain-specific *Hsf1* knockout model (i). **(b), (c), (d), (j), (k), (l)** Bar plots showing the total time (minutes) spent in wake (b,j), NREM sleep (c,k) or REM sleep (d,l) in each phase (Total: ZT0-24, Light: ZT0-12 or Dark: ZT12-24) during SD period for the systemic *Hs1* knockout model (b,c,d) or brain-specific *Hsf1* knockout model (j,k,l). **(e), (m)** Line plots showing the hourly average of the delta power density in NREM sleep (%) during Recovery period for the systemic *Hs1* knockout model (e) or brain-specific *Hsf1* knockout model (m). **(f), (g), (h), (n), (o), (p)** Bar plots showing the total time (minutes) spent in wake (f,n), NREM sleep (g,o) or REM sleep (h,p) in each phase during Recovery period for the systemic *Hs1* knockout model (f,g,h) or brain-specific *Hsf1* knockout model (n,o,p). Data are mean ±s.e.m. Statistical significance was determined using two-way ANOVA with Tukey’s HSD (b,c,d,e,f,g,h,j,k,l,m,n,o,p). *: p<0.05, **: p<0.01. P values for main effect of genotype factor: control or *Hsf1* knockout (Genotype), and interaction effect between genotype and time factors: every hour from ZT1 to ZT24 (Time:Genotype) (e,m).

**Supplementary Figure 6:**
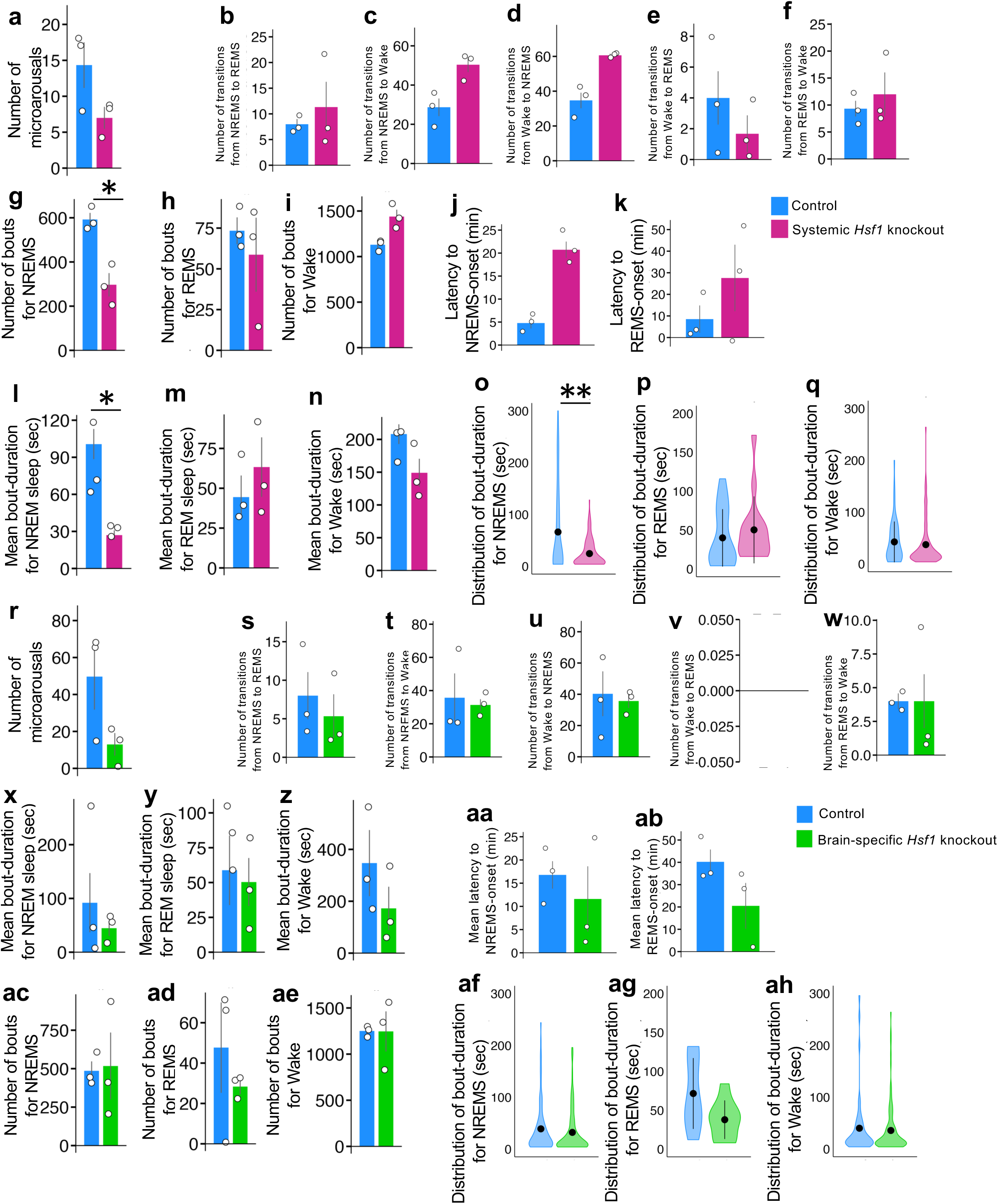
Sleep spectral analysis of systemic and brain specific *Hsf1* knockout mice following sleep deprivation. Results for the systemic (a-q) or brain-specific (r-ah) *Hsf1* knockout model during 2 hours after sleep deprivation (ZT6-8). n=3 per group. **(a), (r)** Bar plots showing the average number of microarousals during the period. **(b), (c), (d), (e), (f), (s), (t), (u), (v), (w)** Bar plots showing the average number of sleep stage transitions from NREM sleep to REM sleep (b,s), NREM sleep to Wake (c,t), Wake to NREM sleep (d,u), Wake to REM sleep (e,v), or REM sleep to Wake (f,w) during the period. **(g), (h), (i), (x), (y), (z)** Bar plots showing the average number of bouts for NREM sleep (g,x), REM sleep (h,y), or Wake (I,z) during the period. **(j), (k), (aa), (ab)** Bar plots showing the mean latencies to the onset of NREM sleep (j,aa), or REM sleep (k,ab) during the period. **(l), (m), (n), (ac), (ad), (ae)** Bar plots showing the average bout-durations (minutes) for NREM sleep(l,ac), REM sleep (m,ad), or Wake (n,ae) during the period. **(o), (p), (q), (af), (ag), (ah)** Violin plots showing distributions of the mean bout durations for NREM sleep (o,af), REM sleep (p,ag), or Wake (q,ah) during the period. Data are mean ±s.e.m. (a-n,r-ae), or median with IQR (o,p,q,af,ag,ah). Statistical significance was determined using two-tailed t-test (a-n,r-ae), or Wilcoxon rank-sum test (o,p,q,af,ag,ah). *: p<0.05, **: p<0.01.

**Supplementary Figure 7:**
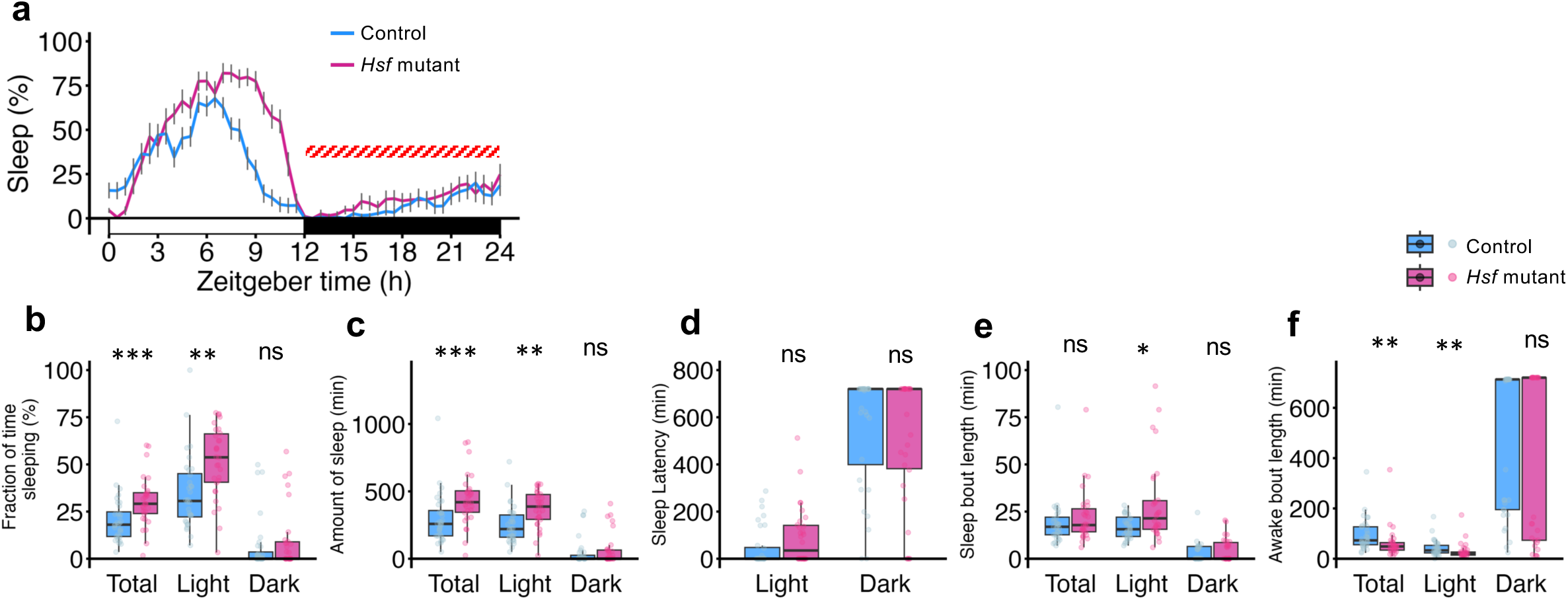
Behavior of *Hsf* mutant flies during SD period. **(a)** Line plot showing the flies’ average proportion of time spent sleeping in consecutive 30min segments (%) during SD period. **(b)** Box plot showing distributions of the mean proportion of time spent sleeping in consecutive 30 minutes segments (%) in Total (ZT0-24), Light (ZT0-12) or Dark (ZT12-24) phase. **(c)** Box plot showing distributions of the mean sleep amount (minutes) in each phase. **(d)** Box plot showing distributions of the mean sleep latency to the first sleep event (minutes) from ZT0 (Light) or ZT12 (Dark). **(e)** Box plot showing distributions of the mean bout-length (minutes) for sleep in each phase. **(f)** Box plot showing distributions of the average bout-length (minutes) for awake in each phase. Data are mean ±s.e.m. (a), or median with IQR and data points for individual average (b,c,d,e,f). Statistical significance was determined using two-step approach (b,c,d,e,f): First, data for each group (control or *Hsf* mutant) and phase (Total, Light or Dark) were assessed for normality using the Shapiro-Wilk test. If the data were normally distributed (p ≥ 0.05), an independent samples t-test was employed to compare the means between independent groups within each phase. If the data significantly deviated from normality (p < 0.05), the Mann-Whitney U test, a non-parametric alternative, was used. All tests were conducted separately for different phases and conditions. *: p<0.05, **: p<0.01, ***: p<0.001, ns: not significant.

